# Daily mitochondrial dynamics in cone photoreceptors

**DOI:** 10.1101/2020.04.10.030262

**Authors:** Michelle M. Giarmarco, Daniel C. Brock, Brian M. Robbings, Whitney M. Cleghorn, Kristine A. Tsantilas, Kellie C. Kuch, William Ge, Kaitlyn M. Rutter, Edward D. Parker, James B. Hurley, Susan E. Brockerhoff

## Abstract

Cone photoreceptors in the retina are exposed to intense daylight, and have higher energy demands in darkness. Cones produce energy using a large cluster of mitochondria. Mitochondria are susceptible to oxidative damage, and healthy mitochondrial populations are maintained by regular turnover. Daily cycles of light exposure and energy consumption suggest that mitochondrial turnover is important for cone health. We investigated the 3-D ultrastructure and metabolic function of zebrafish cone mitochondria throughout the day. At night cones undergo a mitochondrial biogenesis event, corresponding to an increase in the number of smaller, simpler mitochondria and increased metabolic activity. In the daytime, ER and autophagosomes associate more with mitochondria, and mitochondrial size distribution across the cluster changes. We also report dense material shared between cone mitochondria that is extruded from the cell in darkness, sometimes forming extracellular structures. Our findings reveal an elaborate set of daily changes to cone mitochondrial structure and function.

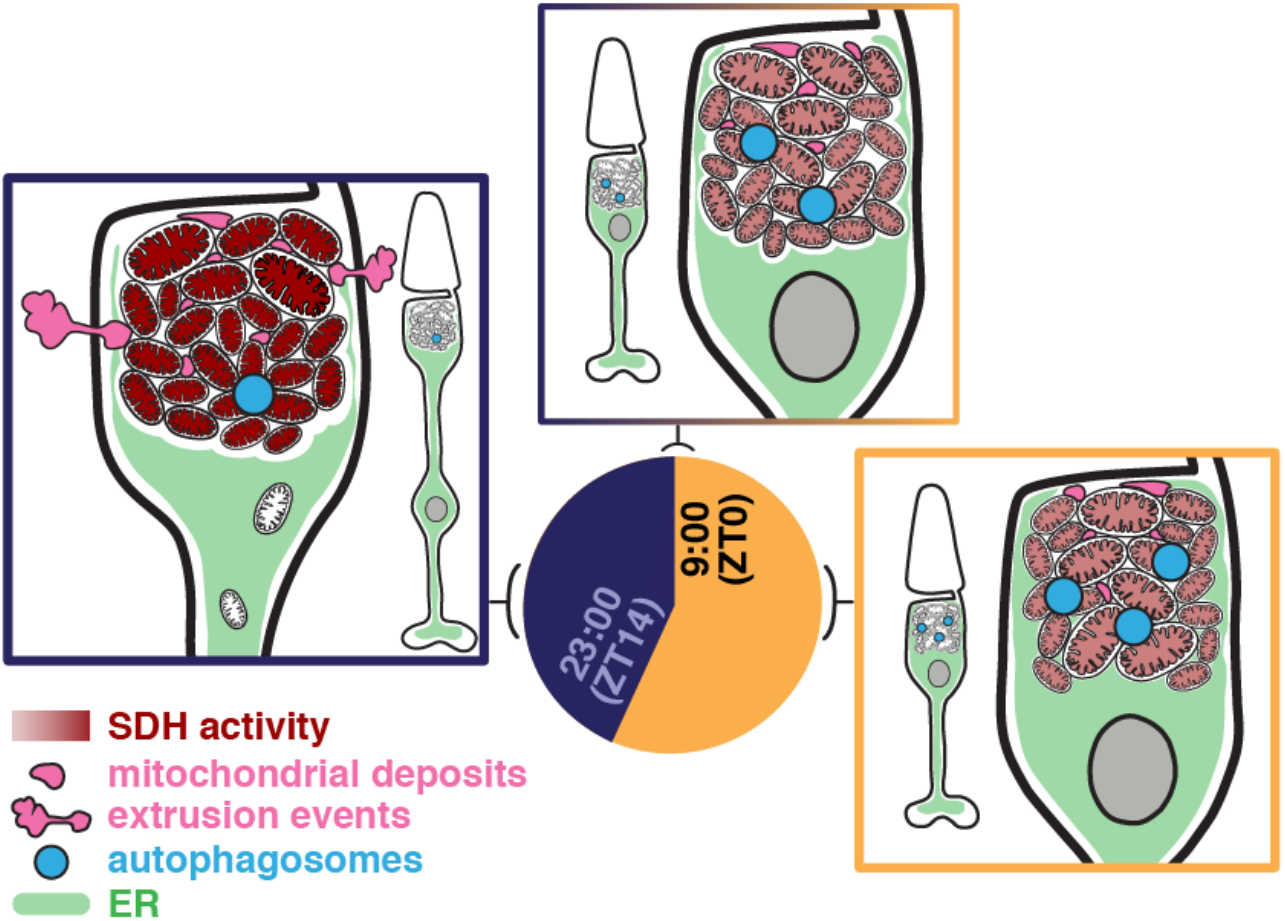

## Introduction

Photoreceptor cells in the retina are highly metabolically active. Their energy demands change throughout the day to support phototransduction (Okawa et al. 2008) and regeneration of outer segment (OS) disks (LaVail 1976). Photoreceptors consume several fold more energy as ATP in darkness than in light (Okawa et al. 2008), and some additional ATP comes from mitochondrial metabolism (Du et al. 2016).

Energy production in most cells is influenced by mitochondrial fission, fusion and new growth (biogenesis). Smaller, fragmented mitochondria typically produce less ATP (Chen et al. 2005). Mitochondrial biogenesis is regulated by many factors including circadian rhythms (de Goede et al. 2018). In neurons it can occur far away from the cell body (Van Laar et al. 2018). Mitochondria can form networks (Bleck et al. 2018) and folds of cristae within mitochondria can be remodeled (Cogliati et al. 2016). These dynamic processes are essential for cell health; mitochondrial dysfunction often accompanies neurodegenerative diseases (Wong et al. 2019; Cieri et al. 2017; Chen & Chan 2009) including retinal degeneration (Litts et al. 2015; Lefevere et al. 2017).

Over 90% of glucose taken up by photoreceptors is used for aerobic glycolysis (Ames et al. 1992; Winkler 1981). Nevertheless, they have a large cluster of mitochondria in the apical portion of the inner segment, the ellipsoid, just below the OS. The density and organization of mitochondrial clusters vary among species, but they are present in photoreceptors of all vertebrates examined, including fish (Tarboush et al. 2014), ground squirrels (Sajdak et al. 2019), mice (Kanow et al. 2017), and humans (Nag & Wadhwa 2016). In some species, all mitochondria reside within the cluster, while mammals with vascularized inner retinas also contain mitochondria at photoreceptor synapses (Stone et al. 2008; Bentmann et al. 2005). In cultured chicken retinas, mitochondrial dynamics are circadian (Chang et al. 2018), but the diurnal changes in mitochondrial structure and function in photoreceptors in intact eyes has not been explored.

In this report we describe daily changes that occur in zebrafish cone photoreceptor mitochondria. Zebrafish provide a useful model to dissect mitochondrial dynamics in specific photoreceptor subclasses. Zebrafish undergo typical vertebrate behavioral and biological circadian rhythms (Vatine et al. 2011; Cahill 2002), and their retinas have four cone subtypes (red, green, blue, and ultraviolet (UV)) organized in a tiered, mosaic pattern. Each cone type can be identified by its unique morphology and position in the outer retina (Raymond et al. 1995), and each maintains a large cluster of mitochondria just below the OS (Tarboush et al. 2014). Our results indicate that mitochondrial clusters in cones are highly dynamic, undergoing diurnal remodeling consistent with enhanced energy production in darkness.

## Results

### Zebrafish cones have more small mitochondria at night

To examine mitochondrial cluster dynamics throughout the day, we collected retinas at 6 timepoints from adult zebrafish under 14h/10h light-dark (LD) or 24 h dark (DD) conditions. Retinas were used for a combination of imaging and biochemical experiments to determine cone mitochondrial form and function. Figure 1A illustrates individual cone cell structures with immunohistochemistry (IHC) and electron microscopy (EM). Cone types were differentiated according to double cone position, nuclear morphology, and presence of a distinct large mitochondrion at the base of UV cone clusters (Figure 1A, white arrows). At night cones extend distally into the retinal pigment epithelium (RPE) by daily retinomotor movements (Figure S2A), a process regulated by light exposure and the circadian clock (Menger et al. 2005; Hodel et al. 2006).

**Figure 1.**
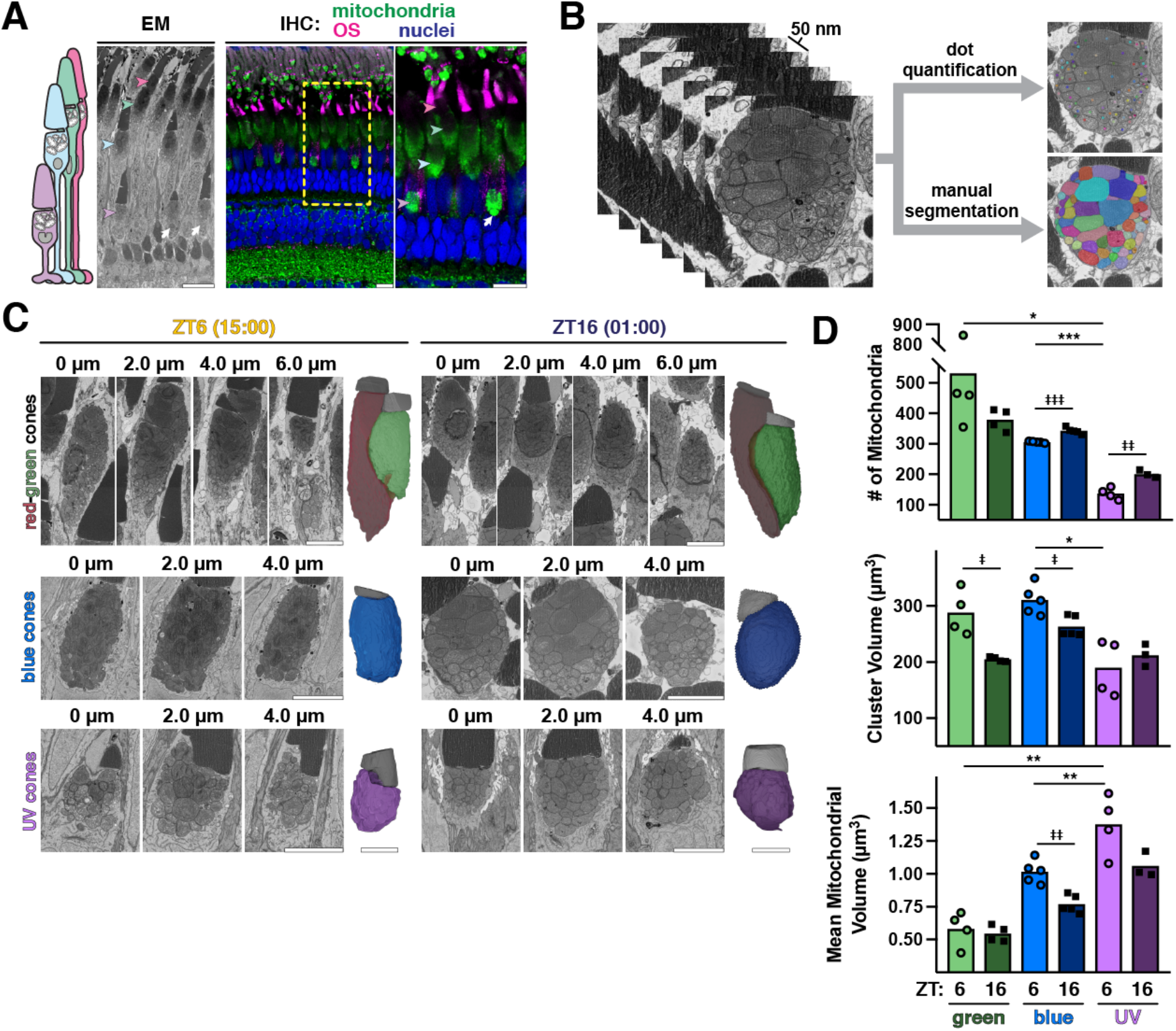
At night cones have more small mitochondria. A) Schematic of zebrafish cone subtypes (left), with EM (middle) and IHC (right) images of zebrafish outer retina. IHC images are stained for all mitochondria (green), red-green cone OS (magenta), and nuclei (blue). Arrowheads indicate corresponding UV, blue, green, and red cone mitochondrial clusters; white arrows, megamitochondria at the UV cluster base. Yellow box indicates zoomed-in area; scale bars, 10 μm. B) Example 50 nm Z-stack from SBFSEM used for 3-D analysis via manual segmentation or rapid dot quantification. C) Z-stacks from SBFSEM with 3-D rendered mitochondrial clusters (colored) and OS (grey) from red-green, blue and UV cones in daytime at ZT6 (15:00) and night at ZT16 (01:00). Scale bars, 5 μm. D) Quantifications of mean mitochondrial number, cluster volume, and mitochondrial volume from dot quantification. Green, blue, and UV cones are represented by respective colors at ZT6 (open circles) and ZT16 (black squares). Table 2 lists p-values and Ns from each group.

We performed detailed 3-D analyses of cone mitochondrial clusters using 50 nm Z-sections collected by serial block-face scanning electron microscopy (SBFSEM) (Figure 1B, Video 1). Image stacks were analyzed using either a rapid dot quantification method or manual segmentation to precisely compare mitochondrial number, size, shape, and location between day (15:00, ZT6) and night (01:00, ZT16). Examples of individual red-green, blue, and UV cone image stacks with corresponding 3-D renderings of mitochondrial clusters and outer segments (OS) at ZT6 and ZT16 are presented in Figure 1C. Figure S1 presents 3-D renderings with mitochondrial numbers and cluster volumes for all cells used in this study.

When analyzed via dot quantification we found that at night, blue and UV cone mitochondria increase in number by 11 ± 1% and 32 ± 4% respectively (Figure 1D, top). These changes did not coincide with an increase in cluster volume; on average, volumes of individual blue and UV cone mitochondria decreased at night by 32 ± 3% and 30 ± 5% respectively (Figure 1D, bottom). While at night single cone clusters contain more small mitochondria, green cones had fewer mitochondria and smaller clusters at night (Figure 1D). This suggests that single and double cone subtypes may undergo different cycles of mitochondrial dynamics.

Mitochondrial cluster shape was also examined for each cone subtype using transgenic zebrafish expressing YFP targeted to cone mitochondria (gnat2:mito-cpYFP) (Giarmarco et al. 2017), counterstained with a variety of antibodies targeting the mitochondrial respiratory chain (Figure S2A; Table 1 lists antibodies used). Individual cone mitochondrial cluster lengths and circularity ratios were calculated, and revealed that for all cone types, cluster length increases at night by ~50% (Figure S2B, top), but cluster width, reflected by the circularity ratio, decreases at this time (Figure S2B, bottom). Light exposure had subtle effects on cluster morphology; the DD group was significantly more elongated in the daytime than the LD group. Both groups exhibited cyclical changes in cluster morphology, indicating regulation by the circadian clock.

### Cone mitochondria within clusters vary in size and complexity

To examine individual mitochondrial morphology, manual segmentation was performed for a subset of cells imaged with SBFSEM (Figure 1B, Video 1). Surface area and volume of each mitochondrion was used to calculate mitochondrial complexity index (MCI), a size-insensitive measure of morphological complexity (Vincent et al. 2019). Mitochondria within clusters were heterogeneous in volume and MCI (Figure 2A), and MCI was poorly correlated with volume (Figure 2B). Four individual mitochondria with the same volume but different MCIs are presented in Figure 2C. Compared to mouse cones, zebrafish cone mitochondria are simpler on average but occupy a larger volume (Figure S3).

**Figure 2.**
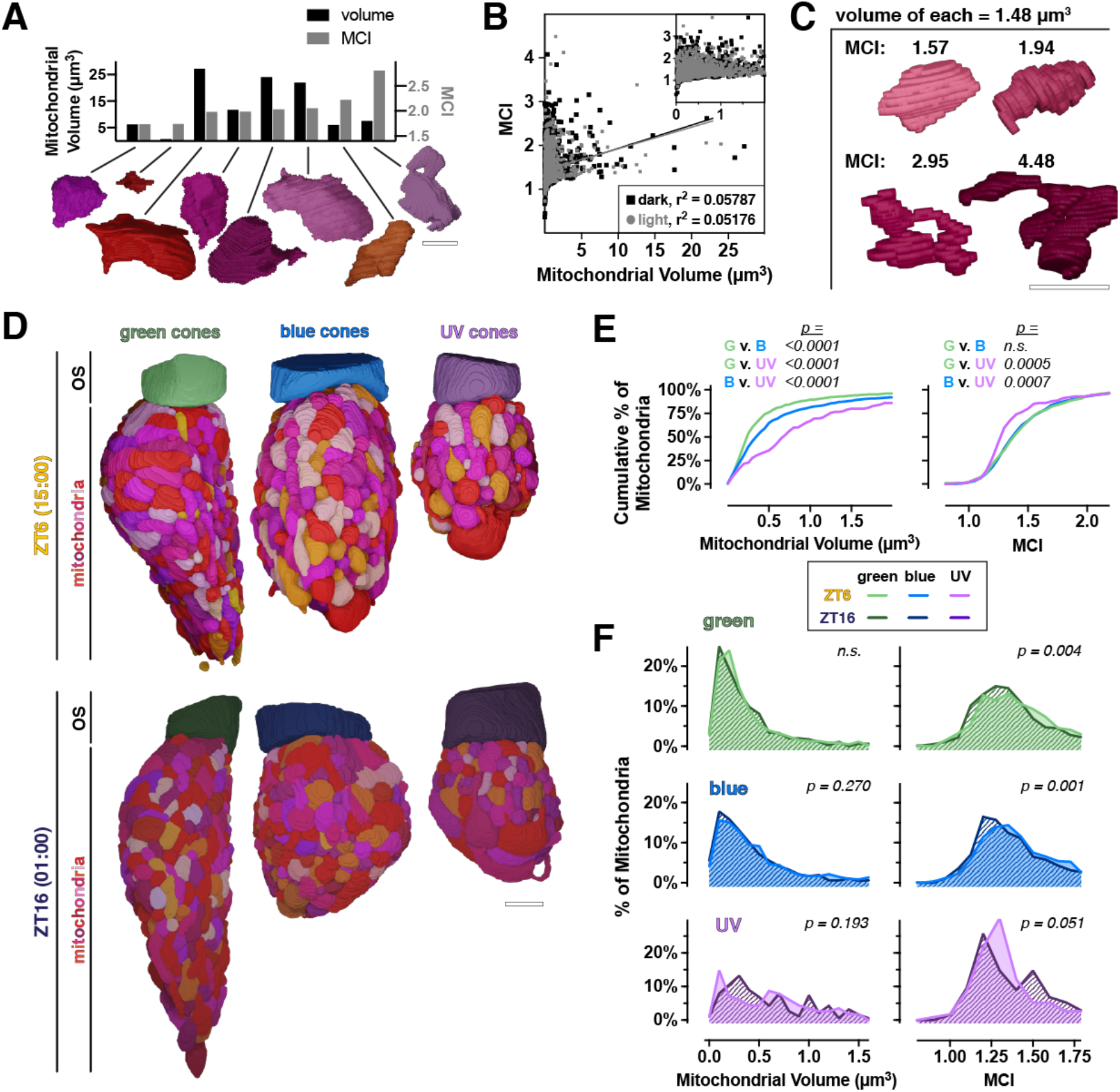
At night cones have more simple mitochondria. A) 3-D renderings of 8 individual cone mitochondria with corresponding quantifications of volume (left Y axis, black bars) and MCI (right Y axis, grey bars). Scale bar, 2 μm. B) Cross correlation plot of mitochondrial volume and MCI for individual mitochondria at ZT16 (black squares) and ZT6 (grey circles). C) 3-D renderings of single mitochondria with equivalent volumes (1.48 μm^3^) over a range of MCIs. Scale bar, 2 μm. D) 3-D renderings from manual segmentation of mitochondria and outer segments (OS) in green, blue and UV cones at ZT6 (top) and ZT16 (bottom). Scale bar, 5 μm. E) Cumulative frequency distributions for mitochondrial volume and MCI comparing green, blue and UV cone mitochondria at ZT6. F) Cumulative frequency distributions for green (top), blue (middle) and UV (bottom) cones, comparing mitochondrial volume and MCI at ZT6 (light lines) and ZT16 (dark lines). ZT6 curves are also presented in (E); Table 2 lists Ns from all groups.

### Mitochondria in green, blue and UV cones are morphologically distinct, and more simple mitochondria appear at night

Our analyses also revealed morphological distinctions between cone subtypes. Clusters in longer-wavelength cones are longer (Figures S1, S2C) and have more mitochondria (Figure 1D). Figure 2D and Video 2 depict 3-D renderings of manually segmented green, blue, and UV cone clusters. At ZT6, green, blue, and UV mitochondria have significantly different distributions of mitochondrial volume, with mitochondrial volumes largest in UV cones and smallest in green cones (Figure 2E, left). Green and blue cone mitochondria have similar MCI distributions, but UV cones have significantly more simple mitochondria (Figure 2E, right).

To compare mitochondrial volume and MCI in cone subtypes between day and night, we analyzed manually-segmented green, blue, and UV cones at ZT6 and ZT16. While volume distributions do not change significantly between day and night for the cells analyzed, the MCI of green and blue cones shifts significantly toward having more simple mitochondria at night. These results suggest that each cone subtype undergoes unique daily regulation to control mitochondrial size and morphology.

### Megamitochondria associate in the cluster core

Cones of several species (Knabe et al. 1997; Utsumi et al. 2020; Tyrrell et al. 2019) including zebrafish (Kim et al. 2005) contain extremely large megamitochondria. To examine distribution of megamitochondria in cone clusters, we 3-D rendered the three largest mitochondria in each cone from Figure 2D in their respective clusters. In all cone subtypes, megamitochondria localize within the cluster core, often in direct contact with one another (Figure 3A). We did not detect striking differences in megamitochondria between day and night with this small sample size, but note that the volume, location and complexity of megamitochondria vary widely between cone subtypes (Figure S4). The largest mitochondria in green cones are relatively uniform in size and of average complexity, while blue and UV cones contained distinct megamitochondria whose volume comprises 12-22% of the entire cluster. We observed several megamitochondria with thin projections extending toward the cluster periphery, and UV cones megamitochondria are unusually complex.

**Figure 3.**
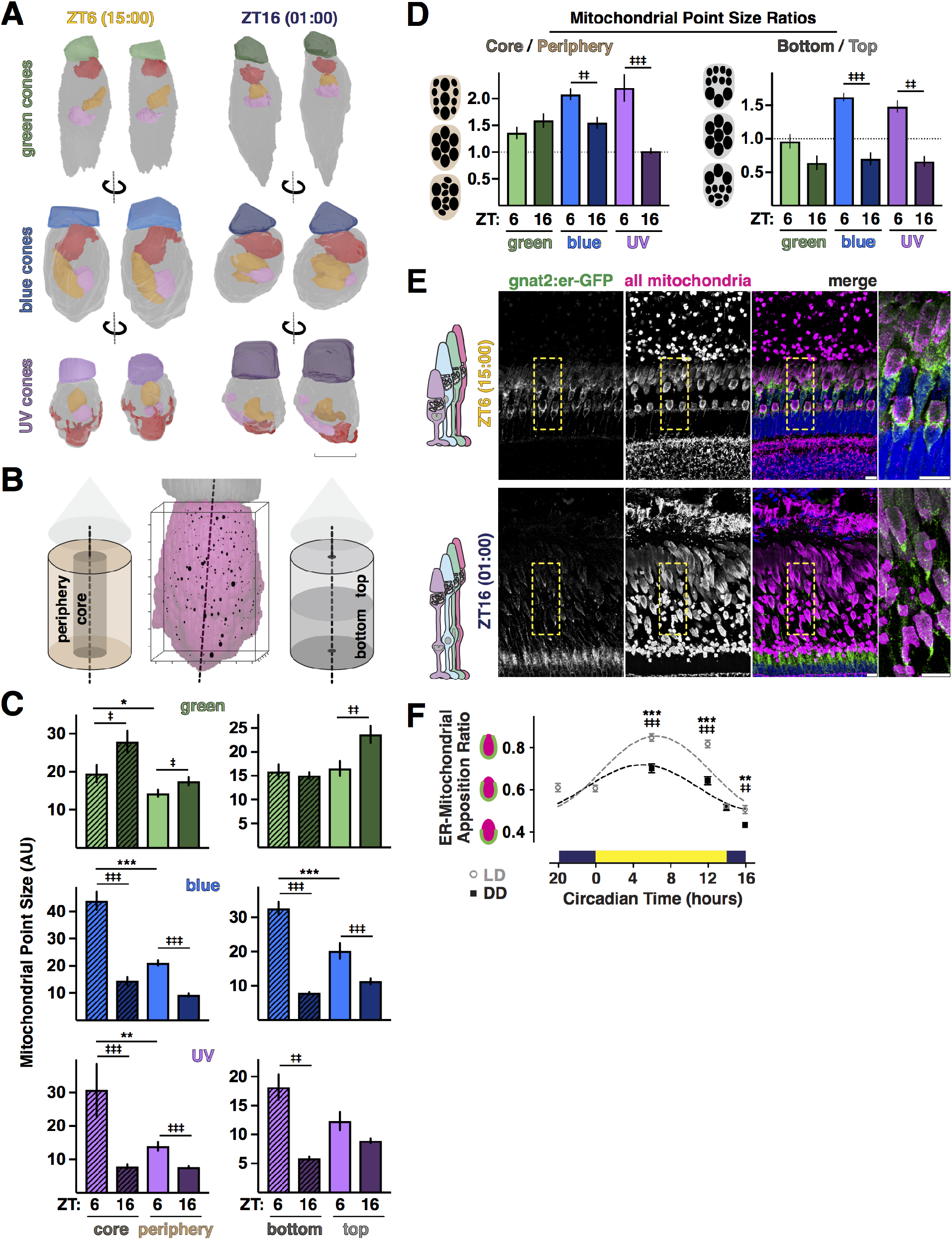
Distribution of mitochondrial size across the cluster changes throughout the day. A) 3-D renderings from manual segmentation of the 3 largest mitochondria in green (top), blue (middle) and UV (bottom) cones from Figure 2D at ZT6 and ZT16. OS, colored respectively; ellipsoids, grey. Largest mitochondrion, red; middle, orange and smallest, pink. Scale bar, 5 μm. B) 3-D rendering of cone ellipsoid (magenta) and OS (grey) overlaid with the corresponding point cloud generated from dot quantification. Individual mitochondria are represented at their X-Y-Z locations in the point cloud, with point size corresponding to relative mitochondrial size. Point clouds were separated into core or peripheral populations, and top or bottom populations relative to the OS. Axis ticks, 1 μm. C) Quantification of mean mitochondrial point size in different regions of the cluster for green (top), blue (middle), and UV (bottom) cones at ZT6 (light colors) and ZT16 (dark colors). D) Mitochondrial point size ratios quantifying regional core-periphery and bottom-top distributions of mitochondrial size reported in (B) for green, blue, and UV cones at ZT6 and ZT16. E) IHC images of transgenic zebrafish outer retina expressing cone-targeted er-GFP (green) overlaid with mitochondrial and nuclear stains (magenta and blue, respectively). Yellow boxes indicate zoomed-in area; scale bars, 10 μm. F) Quantification of mean ER-mitochondrial apposition in blue and UV cones from IHC over time for LD or DD groups. Table 2 lists p-values and Ns from all groups.

### Mitochondria in the cluster core are smaller at night

To more broadly examine the distribution of mitochondrial sizes across the cluster, each mitochondrion from dot quantification was plotted in 3-D using its center X-Y-Z coordinates (Figure 3B, middle). The number of dots needed to track each mitochondrion was reflected by the size of its point in 3-D; larger points represent larger or more branched mitochondria. Points were separated into peripheral and core populations according to distance from a center-axis 3-D line through the cluster (Figure 3B, left). Similarly, points were separated into top and bottom populations according to proximity to the OS or cluster base (Figure 3B, right). Mitochondrial point sizes were then quantified in relation to cluster position (Figure 3C), and ratios of core/periphery and top/bottom were used as measures of relative size distribution (Figure 3D).

At ZT6, blue and UV cones have significantly more large mitochondria at the cluster core, and more small mitochondria at the periphery (Figures 3C and 3D, left). At ZT16 in blue and UV cones, average mitochondrial point size decreases ~70% in the core and ~50% in the periphery; size also decreases ~70% at the bottom and 25-40% at the top of the cluster (Figure 3C). Conceptually, this represents a shift toward more small mitochondria at the base of the cluster at night (Figure 3D). Mitochondrial size distribution in green cones did not significantly change, further evidence for distinct patterns of mitochondrial dynamics between cone subtypes. These data suggest that all cones maintain a population of small mitochondria at the cluster periphery in daytime; at night in single cones, mitochondrial size becomes more uniform across the cluster.

### ER-mitochondrial appositions peak in daytime

In other cells the organelle endoplasmic reticulum (ER) initiates mitochondrial fission (Friedman et al. 2011; Korobova et al. 2013), a process that occurs in the final stages of mitochondria biogenesis (Amiri & Hollenbeck 2008). Cone ER primarily contacts mitochondria at the cluster base and periphery (Giarmarco et al. 2017), where mitochondrial size is most dynamic. We examined the cone ER network surrounding mitochondria using IHC at all timepoints on sections from LD or DD transgenic zebrafish expressing GFP targeted to cone ER (gnat2:er-GFP) (George et al. 2014), and antibodies for the mitochondrial respiratory chain (Figure 3E). To quantify the extent of potential ER contacts for individual blue and UV cone clusters, ER-mitochondrial apposition ratios were calculated by dividing the longest length of ER adjacent to the cluster by the overall cluster length.

In the daytime, ER tightly associates with the entire cluster surface and is densely packed around its base (Figure 3E, top right). At night, apposition ratios decrease as the ER surrounds the cluster more diffusely and extends less toward the OS (Figure 3E, bottom right; Figure 3F). This suggests that the ER network changes at night to accommodate the mitochondrial cluster, and is consistent with the observed shift toward larger mitochondria at the top and periphery of single cone clusters at night (Figure 3D). At most timepoints ER-mitochondrial apposition ratios are lower for the DD group (Figure 3F), suggesting that light exposure drives tighter ER-mitochondrial associations.

### Mitochondrial biogenesis genes peak before night onset

The presence of more mitochondria at night could be due to increased mitochondrial biogenesis. Biogenesis is controlled at the transcriptional level (Scarpulla et al. 2012; Ploumi et al. 2017): upstream cellular signals lead to mitochondrial DNA (mtDNA) replication, protein synthesis and fission creating new mitochondria (Figure 4A). We measured mRNA transcript levels of genes involved with mitochondrial biogenesis using qPCR with whole retinas (Artuso et al. 2012) (Table 3 lists primers used; Table 4 lists fitting parameters for cosinor curves). As a control we examined *aanat2* (M. Wang et al. 2015), which displays robust circadian changes in expression (Figure 4C). The early nuclear transcription factors *pgc1α* and *pgc1β* peak in the daytime, while the mitochondrial transcription factor *tfam,* the mitochondrial DNA polymerase *polg1*, and the mitochondrial deacetylase *sirt3* all peak immediately before night onset (Figure 4B). Together this suggests that canonical mitochondrial biogenesis peaks at night.

**Figure 4.**
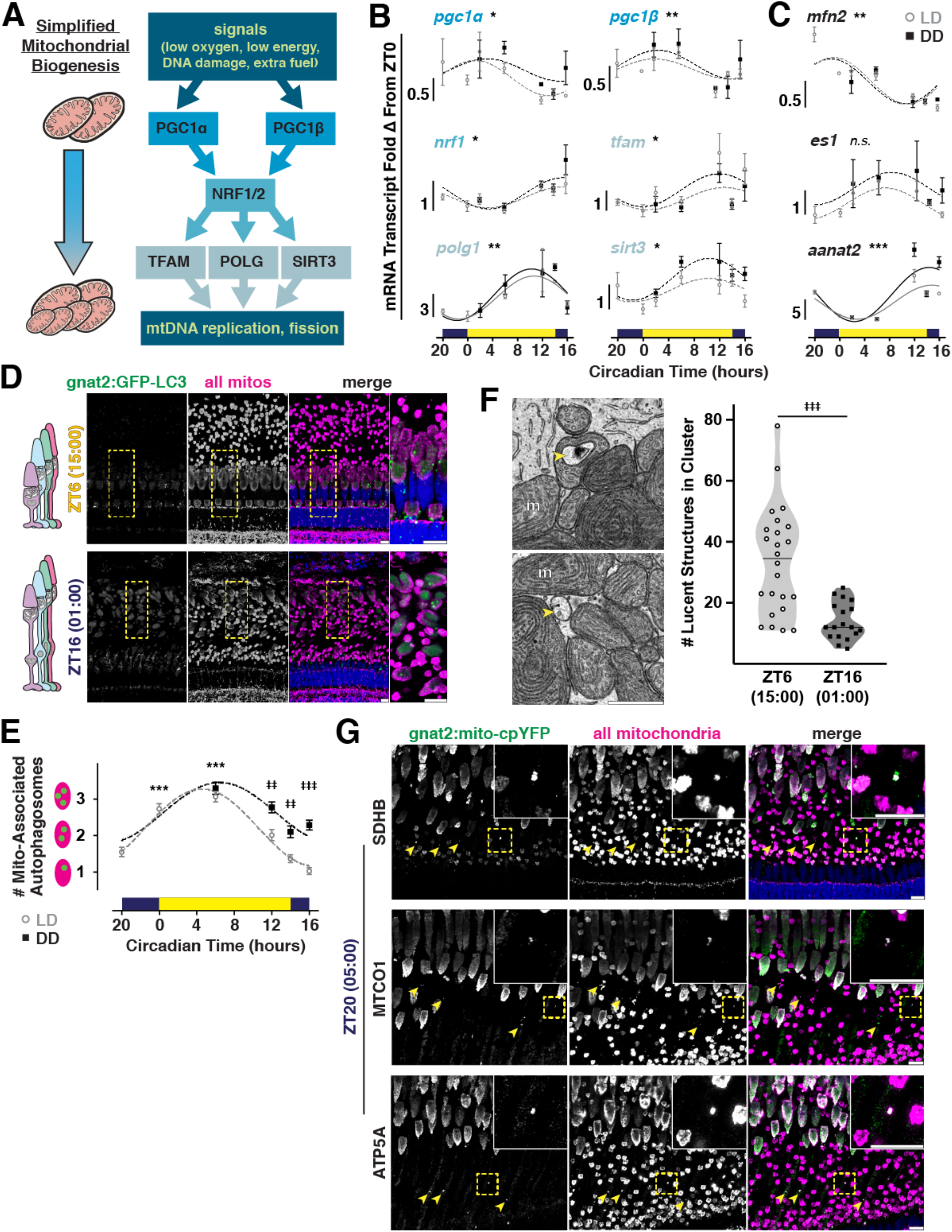
Mitochondrial biogenesis genes peak in the evening, when fewer autophagosomes associate with mitochondrial clusters. A) A simplified pathway for mitochondrial biogenesis. B) and C) Quantification of mRNA transcripts from whole retinas measured using qPCR: in B) 6 mitochondrial biogenesis genes, and in C) the mitochondrial fusion factor *mfn2,* the mitochondrial enlargement factor *es1,* and a control gene *aanat2.* LD (grey open circles), DD (black squares); data normalized to ZT0. D) IHC images of transgenic zebrafish outer retina expressing cone-targeted GFP-LC3 (green) overlaid with mitochondrial and nuclear stains (magenta and blue, respectively). Top row, light (ZT6); bottom row, dark (ZT16). Yellow boxes indicate zoomed-in area; scale bars, 10 μm. E) Quantification of mean mitochondrial LC3-positive puncta in blue and UV cones from IHC over time for LD or DD groups. F) Left, SEM images of lucent autophagosome-like structures (yellow arrowheads) inside and between cone mitochondria (m). Scale bar, 1 μm. Right, violin plots quantifying the number of lucent structures in clusters at ZT6 (empty circles) and ZT16 (black squares); lines represent median. G) IHC images of transgenic cone-targeted mito-cpYFP (green) overlaid with three antibody markers for SDHB, MTCO1, or ATP5A (magenta) and a nuclear stain (blue) at 05:00 (ZT20). Yellow arrowheads and insets indicate mislocalized mitochondria; scale bars, 10 μm. Table 2 lists p-values and Ns from all groups.

Transcripts encoding the mitochondrial fusion protein *mfn2* rise in the morning (Figure 4C, top), suggesting fusion could mediate the corresponding decrease in mitochondrial number. No significant changes were detected in the zebrafish cone mitochondrial enlarging factor *es1* (Masuda et al. 2016) (Figure 4C, middle). Overall we found no striking differences in transcript levels between LD and DD groups, indicating that expression of mitochondrial biogenesis genes is regulated primarily by the circadian clock.

### Mitochondrial-associated autophagosomes peak in daytime

Clearance of damaged or unnecessary mitochondria occurs via a selective form of autophagy called mitophagy (Youle & Narendra 2011). While several pathways can trigger mitophagy (Kawajiri et al. 2010; Strappazzon et al. 2015), all coalesce on recruitment of the cytosolic protein LC3 to maturing mito-autophagosomes. We performed IHC at all timepoints using LD or DD transgenic zebrafish expressing GFP fused to LC3 in cones (gnat2:GFP-LC3) (George et al. 2014), and antibodies for the mitochondrial respiratory chain (Figure 4D). The number of LC3-positive autophagosomes overlapping with blue and UV cone mitochondrial clusters were quantified. The number of mitochondrial-associated autophagosomes increases 2-fold at light onset (Figure 4E). The number of mitochondrial-associated autophagosomes in the DD group remained elevated in the evening, possibly indicative of enhanced mitochondrial turnover in prolonged darkness.

When viewed using EM, autophagosomes appear as lucent, heterogeneous, multivesicular structures, while mitochondria are more electron-dense and contain distinct folds of cristae (Figure 4F, left, yellow arrowheads). We validated our IHC findings by quantifying the number of electron-lucent structures present in cone mitochondrial clusters imaged using SBFSEM. More lucent structures are present in clusters at ZT6 (Figure 4F, right), when the most mitochondrial-associated autophagosomes were detected using IHC. Several clusters contain >30 lucent structures in the daytime, suggestive of an overall mitophagic event at this timepoint.

### Cone mitochondria mislocalize toward the cell body hours before light onset

The fate of degraded mitochondrial material varies by cell type; canonically it enters the endolysosomal pathway (Strappazzon et al. 2015), but in neurons it can translocate and leave the cell (Melentijevic et al. 2017; C.-H. O. Davis et al. 2014). Mitochondria mislocalize toward the nucleus in degenerating human cones and (Litts et al. 2015) and in a zebrafish cone model of mitochondrial calcium overload (Hutto et al. 2020). 4 h before light onset (05:00, ZT20) we observed mislocalized cone mitochondria using IHC with gnat2:mito-cpYFP transgenic zebrafish and 3 different markers of the mitochondrial respiratory chain (Figure 4G, yellow arrowheads). The structures are feasibly the size of a single mitochondrion (0.5-1 μm length), contain both cone mito-cpYFP and respiratory proteins, and lie between the cluster and the nucleus. They were present only at overnight timepoints ZT16 and ZT20, and in multiple animals over 2 generations. While we could not unequivocally identify these structures in SBFSEM, the data suggest that mitochondrial trafficking between the cluster and cell body occurs prior to light onset.

### Mitochondria share material and extrude it from the cell in darkness

In nearly all cones, we found abundant dark deposits inside and between mitochondria at all times of day in SBFSEM images (Figure 5A, top). Similar deposits can be seen in published EM images of cone mitochondrial clusters in other zebrafish (Fig. 2 of Tarboush et al. 2014), walleye (Figs. 5 and 7 of Januschka et al. 1987), frogs (Fig. 2 of Mercurio & Holtzman 1982), pigeons (Fig. 16 of Ishikawa & Yamada 1969), shrews (Figs. 2, 7 and 10 of Lluch et al. 2003), and ground squirrels (Fig. 5 of Sajdak et al. 2019), as well as mice (Figure S3, yellow arrowheads) and albino zebrafish (Figure S5, yellow arrowheads). When manually segmented and 3-D rendered, we found that these electron-dense lamellar whorls can span several micrometers and contact the matrices of multiple mitochondria, as well as the outer segment (Figure 5A, bottom). Most deposits lie inside of single mitochondria, but they appear in the cytosol and occasionally cross the plasma membrane (Figure 5B, top panels). This apparent extrusion most often occurs as single events, but in some cells multiple concurrent exit sites were observed (Figure 5B, bottom panels).

**Figure 5.**
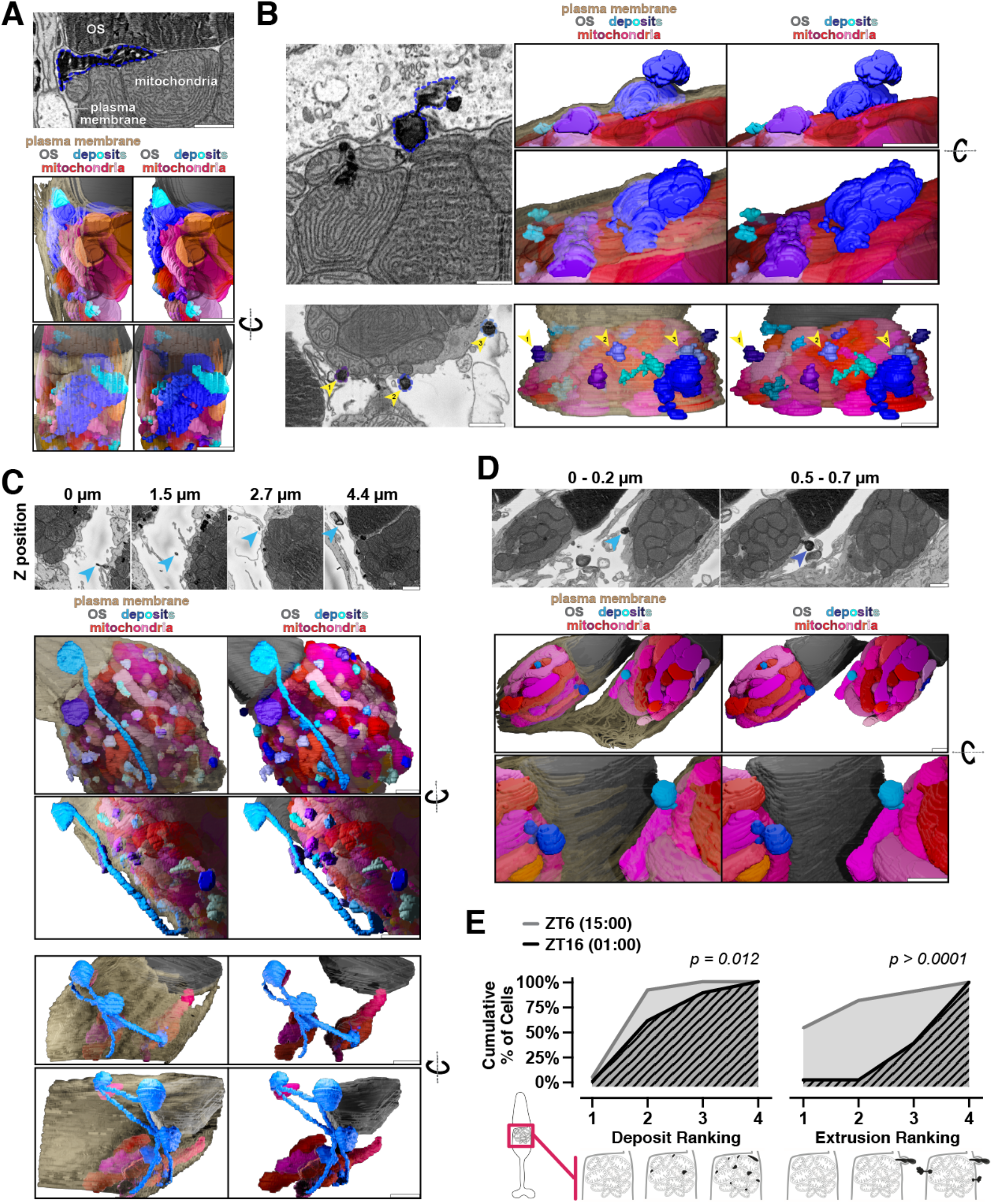
Mitochondria share material and extrude it from the cell in darkness. A) SBFSEM image (top) showing electron-dense deposits associated with cone mitochondria, and corresponding 3-D renderings (bottom). One large deposit (blue outline) below the OS is associated with multiple mitochondria. B) SBFSEM images showing extrusion of mitochondrial-associated deposits from cones, with 3-D renderings. Top, extrusion of one deposit (blue outline). Bottom, multiple extrusion events occurring in one cell (blue-violet outlines, yellow arrows). C) SBFSEM images showing stalks and networks in the extracellular space, and corresponding 3-D renderings. Top, one extruded deposit tethered to mitochondria by a long stalk (blue arrows). Bottom, branched network of extruded material contacting 3 distinct populations of cone mitochondria from one cell. D) Extrusion of mitochondrial-associated deposits from two neighboring rods. Top, SBFSEM minimum intensity projections over 0.2 μm depth highlighting each extrusion event (blue arrows). Bottom, 3-D rendering. All scale bars, 1 μm. Beige, plasma membrane; grey, OS; reds, mitochondria; blues, deposits. E) Quantification of presence of deposits and extrusion events in day (grey, ZT6) and night (black, ZT16). For deposits: (1) no mitochondrial deposits, (2) few mitochondrial deposits, (3) many mitochondrial deposits, (4) every mitochondrion having a deposit. For the number of extrusion events: (1) no events, (2) one event, (3) two or three events, (4) more than three events. Table 2 lists Ns from all groups.

Other neurons can eject degraded mitochondrial material via nanotunnel-like extensions (Melentijevic et al. 2017). 3-D analysis revealed that in cones, the extruded material either remains near the cell surface, or extends away from the cell on stalks that reach as far as 5 μm and terminate in a more diffuse lamellar extracellular sac (Figure 5C, top; Video 3). The stalks are 40-90 nm wide and appear as an electron-dense ring surrounding a hollow core. In the extracellular space stalks and sacs can connect, forming elaborate networks that link discrete populations of mitochondria within a cluster (Figure 5C, bottom; Video 4). These deposits and their extrusion in darkness were observed in nearly all cones but very seldom in rods; in one exceptional instance extrusion was observed in two neighboring rods (Figure 5D).

To determine the extent of cone mitochondrial deposits and extrusion in light and darkness, we used a ranking system to blindly score clusters imaged with SBFSEM.

Deposits were ranked: (1) no mitochondrial deposits to (4) every mitochondrion having a deposit. Similarly the extrusion events were ranked: (1) no events to (4) more than three events. The presence of mitochondrial deposits is only slightly higher at night (Figure 5E, left). However, extrusion events generating stalks and sacs happen exclusively at night, and most cells had multiple events (Figure 5E, right). While the composition of the deposits is not known, they provide a direct physical link between cone mitochondria and the interphotoreceptor matrix.

### Mitochondrial succinate metabolism is more active in darkness due to altered SDH activity

Studies with whole mouse retinas suggest that mitochondrial metabolism is more active in darkness (Du et al. 2016). However ~96% of photoreceptors in mouse retinas are rods (Tsukamoto et al. 2001) while zebrafish retinas have ~50% cones (Raymond et al. 2014; Larison & Bremiller 1990). To measure mitochondrial activity in cones specifically, we performed enzyme histochemistry with fresh frozen retina sections from LD or DD albino zebrafish collected throughout the day. Some albino mouse strains are prone to light-induced retinal degeneration (LaVail et al. 1987), but albino zebrafish used in this study display normal central retinal morphology and mitochondrial clusters (Figure S5). Activities of succinate dehydrogenase (SDH, complex II) and cytochrome c oxidase (COX, complex IV) were assayed separately (Andrews et al. 1999); their roles in mitochondrial metabolism are highlighted in Figure 6A. Histochemistry was performed for all timepoints in parallel, and stain intensities of single cones were measured from light microscopy images (Figure 6B).

**Figure 6.**
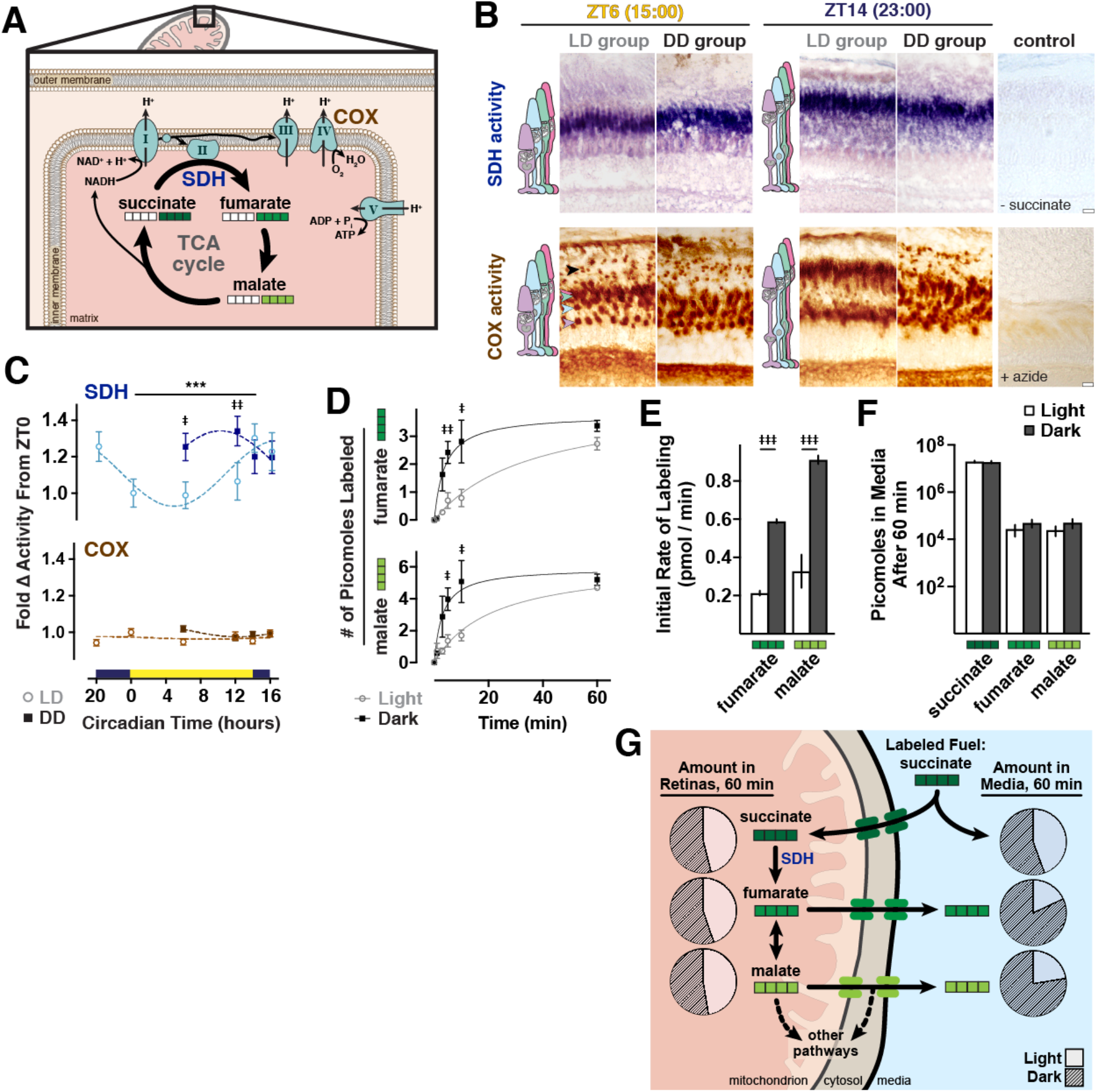
Mitochondrial metabolism is more active in darkness due to altered SDH activity. A) Schematic of SDH in the context of the TCA cycle and electron transport chain, including COX also assayed in (B). White squares represent ^12^C carbons and green squares represent ^13^C carbons for the labeling experiments in (D-F). B) Light microscopy images of histochemical staining for SDH (blue) and COX (brown) activities. Shown are frozen sections from albino zebrafish at ZT6 and ZT14 (23:00) from LD and DD groups. Negative controls lacked substrate or contained an inhibitor. Arrowheads indicate corresponding UV, blue, green, and red cone and rod (black) mitochondrial clusters; scale bars, 10 μm. C) Quantification of SDH and COX stain intensities in single mitochondrial clusters over time in LD (light open circles) or DD (dark squares) groups; data normalized to ZT0. D) Incorporation of ^13^C label from U- ^13^C succinate into fumarate and malate. Whole retinas from light- or dark-adapted zebrafish were incubated in 1 mM U-^13^C succinate with 1 mM glucose in light or dark, and ^13^C incorporation was determined with GS/MS. E) Initial rates of formation of fumarate and malate from U-^13^C succinate in light and dark, determined in the first 5 min of incubation. F) Amounts of U-^13^C labeled succinate, fumarate and malate in media after 60 min light or dark incubation with retinas in 1 mM U-^13^C succinate and 1 mM glucose. Table 2 lists p-values and Ns from all groups. G) Schematic of steadystate metabolites inside and outside of retinas after 60 min incubation with labeled succinate. Pie charts denote relative amounts of succinate, fumarate, and malate in each compartment in light (unfilled areas) and dark (filled areas).

While COX activity remains robust and stable throughout the day, SDH activity in cones increases 25-30% in darkness (Figure 6C). SDH activity in the DD group is elevated at all times of day, suggesting possible repression of SDH activity by light in cones. Additionally SDH activity appears primarily in red-green double cones, compared to strong COX activity observed in all cone subtypes, rods, and the inner retina (Figure 6B, arrowheads).

As further validation, we used metabolite labeling coupled with gas chromatography mass spectroscopy (GC/MS) to assay activity of SDH in light- or dark-adapted zebrafish retinas around ZT6. Whole retinas were incubated with U- ^13^C labeled succinate (dark green boxes, Figure 6A), which SDH converts to U-^13^C fumarate; fumarate is readily interconverted to malate via the enzyme fumarase (Andersen 1980). Labeled fumarate and subsequently labeled malate accumulate faster in dark-adapted retinas (Figure 6D). A 2.75-fold higher initial rate of formation for both fumarate and malate (Figure 6E) is consistent with the higher SDH activity in darkness observed using histochemistry. To determine if succinate uptake was altered in darkness, we also assayed labeled metabolites in the media after 60 min, but did not find significantly lower amounts of labeled succinate (Figure 6F). Figure 6G depicts steady-state metabolite labeling in retinas and media after 60 min in light or darkness; succinate levels are similar in light and dark, but metabolites downstream of SDH accumulate in the media.

## Discussion

### Comparisons of Cone Mitochondria Between Species and Cone Subtypes

There are hundreds of mitochondria in zebrafish cones packed tightly between the nucleus and outer segment. Previous studies revealed that these mitochondria are heterogeneous (Kim et al. 2005; Tarboush et al. 2014; Masuda et al. 2016). In this report we quantified the differences in mitochondrial number, size, complexity and 3-D distribution in zebrafish cone subtypes. Compared with mouse cones (Sloat et al. 2016 and Figure S3), zebrafish cone mitochondria are more numerous, smaller and more densely-packed, with elaborate cristae patterning (Perkins et al. 2003). The dense packing and disparate numbers of mitochondria between cone subtypes may reflect nutrient and oxygen access in the avascular zebrafish retina (Stone et al. 2008); long-wavelength cones could fuel more mitochondria because they are closer to the RPE and blood supply. These spatial constraints, the high SDH activity in long-wavelength cones, and previous studies showing unique metabolic activity in short-wavelength cones (Kam et al. 2019; Nork et al. 1990) suggest that diverse metabolic strategies are needed to support the specific needs of each cone type.

### Mitochondrial Size in Photoreceptors

Mitochondrial size is linked to energetic output; in other cells smaller mitochondria respire less (Bach et al. 2003; Maryanovich et al. 2015; Chen et al. 2005). However, in zebrafish cones it may be the small, morphologically simple mitochondria that contribute more toward energy production. Zebrafish cone mitochondria are smaller and appear simpler at night, when energy demands and mitochondrial respiration are highest. Small mitochondria in cones localize primarily to the cluster periphery, where they have direct access to oxygen and fuels. Cristae in these small mitochondria generally display the organized, linear structure that is linked to higher energetic output (Perkins et al. 2003; Cogliati et al. 2016).

Like cones in other species (T. Ishikawa & Yamada 1969; Knabe & Kuhn 1996) zebrafish cones form megamitochondria, maintained by the enlargement factor ES1 (Masuda et al. 2016; Utsumi et al. 2020). While we did not detect daily changes to *es1* mRNA transcripts in whole retinas, we observed a small population of juxtaposed megamitochondria in all cones throughout the day. The biological role of megamitochondria is not known, but their densely packed cristae suggest they may not perform only canonical respiration (Slautterback 1965). In other tissues, membrane lipids concentrated in cristae stacks of megamitochondria can form a conduit for oxygen (Desaulniers et al. 1996; Urschel & O’Brien 2008). Many megamitochondria in our study exhibited a central mitochondrial body with long fingerlike projections extending toward the cluster periphery; these projections toward the ER could serve as a site for fission during mitochondrial biogenesis (Knabe & Kuhn 1996). Further, there is evidence that cone megamitochondria (Lluch et al. 2003; Knabe et al. 1997) and mitochondrial-derived ellipsosomes (MacNichol et al. 1978; Nag & Bhattacharjee 1995) in other species guide light toward the OS. Zebrafish visual sensitivity is regulated by light and the circadian clock (L. Li & Dowling 1998), so the daily mitochondrial rearrangement we observed may contribute to visual function.

### Mitochondrial Turnover in Photoreceptors

Maintenance of healthy mitochondria and mtDNA is crucial, particularly in the retina. mtDNA mutations (Lefevere et al. 2017) and disrupted autophagy (Rodríguez-Muela et al. 2015) have been associated with retinal degeneration, and aging human cones accumulate mtDNA mutations and other mitochondrial abnormalities (Barron et al. 2001; Nag & Wadhwa 2016). In some forms of age-related macular degeneration, the cone mitochondrial cluster remodels and mitochondria translocate toward the nucleus (Litts et al. 2015).

Genes required for mitochondrial biogenesis (Scarpulla et al. 2012) undergo circadian changes in retinal expression in a manner that supports biogenesis at night onset. At ER-mitochondrial contact sites the ER stimulates mitophagy (Hamasaki et al. 2013) by recruiting autophagic machinery (Huang et al. 2018) and initiating mitochondrial fission (Friedman et al. 2011). In cones, ER wraps most densely around the cluster during the day, when clusters contain more autophagosomes and fewer mitochondria. Together this suggests that cones undergo some degree of daily mitochondrial turnover that involves the ER.

### Succinate metabolism in the retina

Fewer mitochondria during the day could result in reduced SDH activity, which we detected in single cells using histochemistry. Zebrafish cones have higher SDH activity than rods and other retinal cells, consistent with studies of human (Andrews et al. 1999; Cogan & Kuwabara 1959; Barron et al. 2001) and salamander (Moore & Guberg 1974) retinas. Cone SDH activity increases at night, and regulation from the zebrafish circadian clock (P. Li et al. 2008; Cahill 2002) may promote mitochondrial uptake of succinate and malate at night (Cai et al. 2019). However we did not find evidence of increased succinate uptake in darkness, and frozen sections used for histochemistry don’t require transporters for substrate uptake. Thus, our observations of cone SDH activity likely resulted from changes to expression or posttranslational modifications, rather than succinate availability. These experiments tested canonical SDH activity using succinate, though a recent report found evidence of reverse SDH activity using fumarate in mouse retinas (Bisbach et al. 2020).

### Potential modes of SDH regulation

Several species can alter the expression and activity of SDH depending on light exposure (Gu et al. 2002; Davis & Merrett 1974; Popov et al. 2010) and time of day (Akimoto et al. 2005; Reddy et al. 2006). Additionally, metabolites can competitively inhibit SDH (Ackrell et al. 1974; Potter & Dubois 1943), and SDH can be phosphorylated (Salvi et al. 2007; Acín-Pérez et al. 2014; Nath et al. 2015), acetylated (Cimen et al. 2010), and succinylated (Park et al. 2013). mRNA transcripts for the mitochondrial deacetylase SIRT3 increase in retinas prior to night onset, but regulation of SDH by SIRT3 in retinas has not been explored. In photoreceptors both SIRT3 and the mitochondrial desuccinylase SIRT5 are necessary for normal function (Lin et al. 2016). Further, cristae structure can influence respiratory efficiency (Cogliati et al. 2013; Guo et al. 2018), in part by driving SDH supercomplexation (Liu et al. 2018). Changes to the elaborate cristae patterns in cone mitochondria (Perkins et al. 2003 and this study) could direct SDH supercomplexation and energetic output.

### Mitochondrial deposits in photoreceptors

In photoreceptors, darkly stained whorled deposits contact the matrices of multiple mitochondria. The deposits are distinct from melanosomes; they are present in albino zebrafish photoreceptors. While we did not investigate their composition, they closely resemble osmiophilic structures observed in and around mitochondria of other cells: (1) membranous lamellar whorls in rat brown fat mitochondria, posited to play a role in lipid breakdown and storage (Figure 8 of Blanchette-Mackie & Scow 1983); (2) membrane whorls in mitochondria from avian liver cells stimulated with glucagon (Figure 4 of Tarlow et al. 1977); (3) mitochondrial-derived vesicles observed in cultured mammalian cells that selectively send contents to the lysosomal pathway (Soubannier et al. 2012; Sugiura et al. 2014); (4) peroxisome precursors derived from both mitochondria and ER (Sugiura et al. 2017). Further, it has been reported that stacks of ER can encircle mitochondria, forming a lamellar structure to initiate mitophagy (Huang et al. 2018). Collectively this suggests that photoreceptor mitochondrial deposits may play roles in lipid homeostasis and/or mitochondrial clearance.

### Nightly extrusion of mitochondrial material

Mitochondrial deposits appear to be extruded from cones at night. Photoreceptor inner segments use endocytosis to take up components of the interphotoreceptor matrix (IPM) (Hollyfield, Varner, Rayborn, Liou, et al. 1985; Hollyfield, Varner, Rayborn & Bridges 1985; Hollyfield & Rayborn 1987), but the events we observed are morphologically distinct. They are also distinct from exocytosis occurring at photoreceptor synapses (Rea et al. 2004; Wen et al. 2017), and from the mitochondrial extrusion that occurs during reticulocyte maturation (Gasko & Danon 1972; Simpson & Kling 1968), inflammatory response (Unuma et al. 2015), or following mitoptosis (Lyamzaev et al. 2008; Géminard et al. 2002). Unlike these events, the extrusion of cone mitochondrial deposits lacks obvious fusion with the plasma membrane, and the released material remains at least transiently associated with the cell surface and underlying mitochondria.

The secreted material can make connections to the extracellular space via thin hollow stalks that terminate in diffuse lamellar sacs oriented toward the outer segment. Other reports of neurons releasing and tethering mitochondrial material in exophers (Melentijevic et al. 2017) and evulsions (Davis et al. 2014) report thin stalks ~200 nm wide, and terminal multivesiculate sacs of ~3 μm diameter. The extracellular structures attached to photoreceptors in our study were smaller (~60 nm wide stalks and ~1 μm diameter sacs), and sacs did not contain vesicles. The stalks are also smaller than and distinct from tunneling nanotubes, cytoplasmic bridges that connect neighboring cells to transfer material (Gerdes & Carvalho 2008) including mitochondria (Lu et al. 2017; X. Wang & Gerdes 2015; Sartori-Rupp et al. 2019). RPE cells can use tunneling nanotubes to transfer material (Wittig et al. 2012), and photoreceptors are reported to exchange proteins via an unknown mechanism (Ortin-Martinez et al. 2017), but we did not observe connections between cones.

The stalks and sacs we observed also form extracellular networks with material from the same cell that could link cones to the IPM. The IPM is a complicated scaffold of carbohydrates and bound proteins that supports photoreceptors (Ishikawa et al. 2015; Hollyfield 1999; Hollyfield et al. 1998). Metabolite levels vary in different regions of the IPM (Adler & Southwick 1992), and cones occupy a distinct microenvironment there (Johnson et al. 1986). At night the zebrafish outer retina becomes disorganized, so the extracellular structures we describe in this report could help physically anchor cones to the IPM.

In summary, we report a quantitative summary of daily mitochondrial dynamics in cone photoreceptors. At night, when energy demands are highest, a mitochondrial biogenesis event leads to increases in metabolic activity and the number of small mitochondria. In the day, mitochondrial number decreases, perhaps mediated by fusion and/or mitophagy. We also report a dense lamellar material that is shared between mitochondria and remarkably leaves the cell at night, sometimes forming extracellular networks. Cone mitochondria undergo daily changes that support energy production at night, regulated by light and the circadian clock. Elucidating the makeup of mitochondrial deposits, the regulation of SDH, the role of ER-mitochondrial contacts, and overall rates and locations of mitochondrial turnover in healthy cones will contribute to a wider understanding of how mitochondrial abnormalities in aging and disease affect vision.

## Acknowledgments

Funding for this work was provided by the UW Art and Rita Levinson undergraduate research scholarship (D.C.B.), NIH NEI 5T32-EY007031 (M.M.G.), NIH NEI RO1-EY026020 (S.E.B.), NIH NEI RO1-EY06641 (J.B.H.), NIH NIA T32-AG000057 (K.A.T.), and UW Vision Core grant NIH NEI P30-EY001730 (to Maureen Neitz). We thank Amandeep S. Dhami, Ashlee D. Evans, Carson Adams, Stephanie R. Sloat, and Alexy J. Merz for help with data analysis and thoughtful discussion; Stan Kim provided zebrafish care at the UW ISCRM Aquatics Center.

## Author Contributions

Conceptualization: M.M.G., D.C.B. and S.E.B. Methodology: M.M.G., D.C.B., B.M.R., W.M.C., K.A.T., and W.G. Validation: M.M.G., D.C.B., B.M.R., and W.M.C. Formal analysis: M.M.G. and D.C.B. Investigation: M.M.G., D.C.B., B.M.R., W.M.C., K.A.T., W.G., K.C.K., K.M.R. and E.D.P. Resources: E.D.P. Writing – original draft: M.M.G. Writing – review and editing: D.C.B., J.B.H. and S.E.B. Visualization: M.M.G. Supervision and Project administration: M.M.G. and S.E.B. Funding acquisition: J.B.H. and S.E.B.

## References

Acín-Pérez, R. et al., 2014. ROS-triggered phosphorylation of complex II by Fgr kinase regulates cellular adaptation to fuel use. Cell Metabolism, 19(6), pp.1020–1033.

Ackrell, B.A., Kearney, E.B. & Mayr, M., 1974. Role of oxalacetate in the regulation of mammalian succinate dehydrogenase. The Journal of biological chemistry, 249(7), pp.2021–2027.

Adler, A.J. & Southwick, R.E., 1992. Distribution of glucose and lactate in the interphotoreceptor matrix. Ophthalmic research, 24(4), pp.243–252.

Akimoto, H., Kinumi, T. & Ohmiya, Y., 2005. Circadian rhythm of a TCA cycle enzyme is apparently regulated at the translational level in the dinoflagellate Lingulodinium polyedrum. Journal of Biological Rhythms, 20(6), pp.479–489.

Ames, A. et al., 1992. Energy metabolism of rabbit retina as related to function: high cost of Na+ transport. The Journal of neuroscience: the official journal of the Society for Neuroscience, 12(3), pp.840–853.

Amiri, M. & Hollenbeck, P.J., 2008. Mitochondrial biogenesis in the axons of vertebrate peripheral neurons. Developmental Neurobiology, 68(11), pp.1348–1361.

Andersen, B., 1980. Lack of deviation from Michaelis--Menten kinetics for pig heart fumarase. Biochemical Journal, 189(3), pp.653–654.

Andrews, R.M. et al., 1999. Histochemical localisation of mitochondrial enzyme activity in human optic nerve and retina. British Journal of Ophthalmology, 83(2), pp.231–235.

Antinucci, P. & Hindges, R., 2016. A crystal-clear zebrafish for in vivo imaging. Scientific Reports, 6, p.29490.

Artuso, L. et al., 2012. Mitochondrial DNA metabolism in early development of zebrafish (Danio rerio). Biochimica et biophysica acta, 1817(7), pp.1002–1011.

Bach, D. et al., 2003. Mitofusin-2 determines mitochondrial network architecture and mitochondrial metabolism. A novel regulatory mechanism altered in obesity. The Journal of biological chemistry, 278(19), pp.17190–17197.

Barron, M.J. et al., 2001. Mitochondrial abnormalities in ageing macular photoreceptors. Investigative Ophthalmology & Visual Science, 42(12), pp.3016–3022.

Bentmann, A. et al., 2005. Divergent distribution in vascular and avascular mammalian retinae links neuroglobin to cellular respiration. The Journal of biological chemistry, 280(21), pp.20660–20665.

Bisbach, C. M. et al., 2020. Succinate can shuttle reducing power from the hypoxic retina to the O_2_-rich pigment epithelium. CellReports, in press.

Blanchette-Mackie, E.J. & Scow, R.O., 1983. Movement of lipolytic products to mitochondria in brown adipose tissue of young rats: an electron microscope study. The Journal of Lipid Research, 24(3), pp.229–244.

Bleck, C.K.E. et al., 2018. Subcellular connectomic analyses of energy networks in striated muscle. Nature Communications, 9(1), p.5111.

Cahill, G.M., 2002. Clock mechanisms in zebrafish. Cell and tissue research, 309(1), pp.27–34.

Cai, T. et al., 2019. The circadian protein CLOCK regulates cell metabolism via the mitochondrial carrier SLC25A10. Biochimica et biophysica acta. Molecular cell research, 1866(8), pp.1310–1321.

Chang, J.Y.-A. et al., 2018. Circadian Regulation of Mitochondrial Dynamics in Retinal Photoreceptors. Journal of Biological Rhythms, 33(2), pp.151–165.

Chen, H. & Chan, D.C., 2009. Mitochondrial dynamics-fusion, fission, movement, and mitophagy-in neurodegenerative diseases. Human Molecular Genetics, 18(R2), pp.R169–76.

Chen, H., Chomyn, A. & Chan, D.C., 2005. Disruption of fusion results in mitochondrial heterogeneity and dysfunction. The Journal of biological chemistry, 280(28), pp.26185–26192.

Chinchore, Y. et al., 2017. Glycolytic reliance promotes anabolism in photoreceptors. eLife, 6.

Cieri, D., Brini, M. & Calì, T., 2017. Emerging (and converging) pathways in Parkinson’s disease: keeping mitochondrial wellness. Biochemical and Biophysical Research Communications, 483(4), pp.1020–1030.

Cimen, H. et al., 2010. Regulation of succinate dehydrogenase activity by SIRT3 in mammalian mitochondria. Biochemistry, 49(2), pp.304–311.

Cogan, D.G. & Kuwabara, T., 1959. Tetrazolium Studies on the Retina: II. Substrate Dependent Patterns. The journal of histochemistry and cytochemistry: official journal of the Histochemistry Society, 7(5), pp.334–341.

Cogliati, S. et al., 2013. Mitochondrial cristae shape determines respiratory chain supercomplexes assembly and respiratory efficiency. Cell, 155(1), pp.160–171.

Cogliati, S., Enriquez, J.A. & Scorrano, L., 2016. Mitochondrial Cristae: Where Beauty Meets Functionality. Trends in biochemical sciences, 41(3), pp.261–273.

Davis, B. & Merrett, M.J., 1974. The Effect of Light on the Synthesis of Mitochondrial Enzymes in Division-synchronized Euglena Cultures. Plant physiology, 53(4), pp.575–580.

Davis, C.-H.O. et al., 2014. Transcellular degradation of axonal mitochondria. Proceedings of the National Academy of Sciences, 111(26), pp.9633–9638.

de Goede, P. et al., 2018. Circadian rhythms in mitochondrial respiration. Journal of Molecular Endocrinology, 60(3), pp.R115–R130.

Desaulniers, N., Moerland, T.S. & Sidell, B.D., 1996. High lipid content enhances the rate of oxygen diffusion through fish skeletal muscle. The American journal of physiology, 271(1 Pt 2), pp.R42–7.

Du, J. et al., 2016. Phototransduction influences metabolic flux and nucleotide metabolism in mouse retina. The Journal of biological chemistry, 291(9), pp.4698–4710.

Friedman, J.R. et al., 2011. ER tubules mark sites of mitochondrial division. Science, 334(6054), pp.358–362.

Gasko, O. & Danon, D., 1972. Deterioration and disappearance of mitochondria during reticulocyte maturation. Experimental Cell Research, 75(1), pp.159–169.

George, A.A. et al., 2014. Synaptojanin 1 is required for endolysosomal trafficking of synaptic proteins in cone photoreceptor inner segments. PLoS ONE, 9(1), p.e84394.

Gerdes, H.-H. & Carvalho, R.N., 2008. Intercellular transfer mediated by tunneling nanotubes. Current opinion in cell biology, 20(4), pp.470–475.

Géminard, C., de Gassart, A. & Vidal, M., 2002. Reticulocyte maturation: mitoptosis and exosome release. Biocell: official journal of the Sociedades Latinoamericanas de Microscopia Electronica … et. al, 26(2), pp.205–215.

Giarmarco, M.M. et al., 2017. Mitochondria Maintain Distinct Ca^2+^ Pools in Cone Photoreceptors. Journal of Neuroscience, 37(8), pp.2061–2072.

Gu, F.K. et al., 2002. A comparative study on the electron microscopic enzymo-cytochemistry of Paramecium bursaria from light and dark cultures. European Journal of Protistology, 38(3), pp.267–278.

Guo, R. et al., 2018. Structure and mechanism of mitochondrial electron transport chain. Biomedical journal, 41(1), pp.9–20.

Hamasaki, M. et al., 2013. Autophagosomes form at ER–mitochondria contact sites. Nature, 495(7441), pp.389–393.

Hodel, C., Neuhauss, S.C.F. & Biehlmaier, O., 2006. Time course and development of light adaptation processes in the outer zebrafish retina. The Anatomical Record, 288A(6), pp.653–662.

Hollyfield, J.G., 1999. Hyaluronan and the functional organization of the interphotoreceptor matrix. Investigative Ophthalmology & Visual Science, 40(12), pp.2767–2769.

Hollyfield, J.G. & Rayborn, M.E., 1987. Endocytosis in the inner segment of rod photoreceptors: analysis of Xenopus laevis retinas using horseradish peroxidase. Experimental eye research, 45(5), pp.703–719.

Hollyfield, J.G. et al., 1998. Hyaluronan in the interphotoreceptor matrix of the eye: species differences in content, distribution, ligand binding and degradation. Experimental eye research, 66(2), pp.241–248.

Hollyfield, J.G., Varner, H.H., Rayborn, M.E. & Bridges, C.D., 1985. Participation of photoreceptor cells in retrieval and degradation of components in the interphotoreceptor matrix. Progress in clinical and biological research, 190, pp.171–175.

Hollyfield, J.G., Varner, H.H., Rayborn, M.E., Liou, G.I., et al., 1985. Endocytosis and degradation of interstitial retinol-binding protein: differential capabilities of cells that border the interphotoreceptor matrix. The Journal of cell biology, 100(5), pp.1676–1681.

Huang, Y. et al., 2018. A “Lamellar structure” contributes to autophagosome biogenesis and mitophagy in zebrafish hepatocytes. Fish & shellfish immunology, 81, pp.83–91.

Hutto, R.A. et al., 2020. Increasing Ca^2+^ in photoreceptor mitochondria alters metabolites, accelerates photoresponse recovery, and reveals adaptations to mitochondrial stress. Cell Death and Differentiation, 27(3), pp.1067–1085.

Ishikawa, M., Sawada, Y. & Yoshitomi, T., 2015. Structure and function of the interphotoreceptor matrix surrounding retinal photoreceptor cells. Experimental eye research, 133, pp.3–18.

Ishikawa, T. & Yamada, E., 1969. Atypical mitochondria in the ellipsoid of the photoreceptor cells of vertebrate retinas. Investigative ophthalmology, 8(3), pp.302–316.

Januschka, M.M. et al., 1987. The ultrastructure of cones in the walleye retina. Vision research, 27(3), pp.327–341.

Johnson, L.V., Hageman, G.S. & Blanks, J.C., 1986. Interphotoreceptor matrix domains ensheath vertebrate cone photoreceptor cells. Investigative Ophthalmology & Visual Science, 27(2), pp.129–135.

Kam, J.H. et al., 2019. Mitochondrial absorption of short wavelength light drives primate blue retinal cones into glycolysis which may increase their pace of aging. Visual Neuroscience, 36, p.E007.

Kanow, M.A. et al., 2017. Biochemical adaptations of the retina and retinal pigment epithelium support a metabolic ecosystem in the vertebrate eye. eLife, 6.

Kawajiri, S. et al., 2010. PINK1 is recruited to mitochondria with parkin and associates with LC3 in mitophagy. FEBS Letters, 584(6), pp.1073–1079.

Kim, J. et al., 2005. The presence of megamitochondria in the ellipsoid of photoreceptor inner segment of the zebrafish retina. Anatomia, Histologia, Embryologia: Journal of Veterinary Medicine Series C, 34(6), pp.339–342.

Knabe, W. & Kuhn, H.J., 1996. Morphogenesis of megamitochondria in the retinal cone inner segments of Tupaia belangeri (Scandentia). Cell and tissue research, 285(1), pp.1–9.

Knabe, W., Skatchkov, S. & Kuhn, H.J., 1997. “Lens mitochondria” in the retinal cones of the tree-shrew Tupaia belangeri. Vision research, 37(3), pp.267–271.

Korobova, F., Ramabhadran, V. & Higgs, H.N., 2013. An actin-dependent step in mitochondrial fission mediated by the ER-associated formin INF2. Science, 339(6118), pp.464–467.

Kwan, K.M. et al., 2007. The Tol2kit: A multisite gateway-based construction kit forTol2 transposon transgenesis constructs. Developmental Dynamics, 236(11), pp.3088–3099.

Larison, K.D. & Bremiller, R., 1990. Early onset of phenotype and cell patterning in the embryonic zebrafish retina. Development, 109(3), pp.567–576.

LaVail, M.M., 1976. Rod outer segment disk shedding in rat retina: relationship to cyclic lighting. Science, 194(4269), pp.1071–1074.

LaVail, M.M. et al., 1987. Genetic regulation of light damage to photoreceptors. Investigative Ophthalmology & Visual Science, 28(7), pp.1043–1048.

Lefevere, E. et al., 2017. Mitochondrial dysfunction underlying outer retinal diseases. Mitochondrion, 36, pp.66–76.

Li, L. & Dowling, J.E., 1998. Zebrafish visual sensitivity is regulated by a circadian clock. Visual Neuroscience, 15(5), pp.851–857.

Li, P. et al., 2008. CLOCK is required for maintaining the circadian rhythms of Opsin mRNA expression in photoreceptor cells. The Journal of biological chemistry, 283(46), pp.31673–31678.

Lin, J.B. et al., 2016. NAMPT-Mediated NAD(+) Biosynthesis Is Essential for Vision In Mice. CellReports, 17(1), pp.69–85.

Litts, K.M. et al., 2015. Inner Segment Remodeling and Mitochondrial Translocation in Cone Photoreceptors in Age-Related Macular Degeneration With Outer Retinal Tubulation. Investigative Opthalmology & Visual Science, 56(4), pp.2243–2253.

Liu, F. et al., 2018. The interactome of intact mitochondria by cross-linking mass spectrometry provides evidence for coexisting respiratory supercomplexes. Molecular & Cellular Proteomics, 17(2), pp.216–232.

Lluch, S., López-Fuster, M.J. & Ventura, J., 2003. Giant mitochondria in the retina cone inner segments of shrews of genus Sorex (Insectivora, Soricidae). The anatomical record. Part A, Discoveries in molecular, cellular, and evolutionary biology, 272(2), pp.484–490.

Lu, J. et al., 2017. Tunneling nanotubes promote intercellular mitochondria transfer followed by increased invasiveness in bladder cancer cells. Oncotarget, 8(9), pp.15539–15552.

Lyamzaev, K.G. et al., 2008. Novel mechanism of elimination of malfunctioning mitochondria (mitoptosis): formation of mitoptotic bodies and extrusion of mitochondrial material from the cell. Biochimica et biophysica acta, 1777(7-8), pp.817–825.

MacNichol, E.F. et al., 1978. Ellipsosomes: organelles containing a cytochrome-like pigment in the retinal cones of certain fishes. Science, 200(4341), pp.549–552.

Maryanovich, M. et al., 2015. An MTCH2 pathway repressing mitochondria metabolism regulates haematopoietic stem cell fate. Nature Communications, 6, p.7901.

Masuda, T., Wada, Y. & Kawamura, S., 2016. ES1 is a mitochondrial enlarging factor contributing to form mega-mitochondria in zebrafish cones. Scientific Reports, 6, p.22360.

Melentijevic, I. et al., 2017. C. elegans neurons jettison protein aggregates and mitochondria under neurotoxic stress. Nature, 542(7641), pp.367–371.

Menger, G.J., Koke, J.R. & Cahill, G.M., 2005. Diurnal and circadian retinomotor movements in zebrafish. Visual Neuroscience, 22(2), pp.203–209.

Mercurio, A.M. & Holtzman, E., 1982. Smooth endoplasmic reticulum and other agranular reticulum in frog retinal photoreceptors. Journal of neurocytology, 11(2), pp.263–293.

Millard, P. et al., 2012. IsoCor: correcting MS data in isotope labeling experiments. Bioinformatics (Oxford, England), 28(9), pp.1294–1296.

Moore, C.L. & Guberg, E.R., 1974. The distribution of succinic semialdehyde dehydrogenase in the brain and retina of the tiger salamander. Brain Research, 67(3), pp.467–478.

Nag, T.C. & Bhattacharjee, J., 1995. Retinal ellipsosomes: morphology, development, identification, and comparison with oil droplets. Cell and tissue research, 279(3), pp.633–637.

Nag, T.C. & Wadhwa, S., 2016. Immunolocalisation pattern of complex I-V in ageing human retina: Correlation with mitochondrial ultrastructure. Mitochondrion, 31, pp.20–32.

Nath, A.K. et al., 2015. PTPMT1 Inhibition Lowers Glucose through Succinate Dehydrogenase Phosphorylation. CellReports, 10(5), pp.694–701.

Nork, T.M. et al., 1990. Distribution of carbonic anhydrase among human photoreceptors. Investigative Ophthalmology & Visual Science, 31(8), pp.1451–1458.

Okawa, H. et al., 2008. ATP consumption by mammalian rod photoreceptors in darkness and in light. Current biology: CB, 18(24), pp.1917–1921.

Ortin-Martinez, A. et al., 2017. A Reinterpretation of Cell Transplantation: GFP Transfer From Donor to Host Photoreceptors. Stem cells (Dayton, Ohio), 35(4), pp.932–939.

Park, J. et al., 2013. SIRT5-mediated lysine desuccinylation impacts diverse metabolic pathways. Molecular Cell, 50(6), pp.919–930.

Perkins, G.A., Ellisman, M.H. & Fox, D.A., 2003. Three-dimensional analysis of mouse rod and cone mitochondrial cristae architecture: bioenergetic and functional implications. Molecular vision, 9, pp.60–73.

Ploumi, C., Daskalaki, I. & Tavernarakis, N., 2017. Mitochondrial biogenesis and clearance: a balancing act. The FEBS journal, 284(2), pp.183–195.

Popov, V.N. et al., 2010. Succinate dehydrogenase in Arabidopsis thaliana is regulated by light via phytochrome A. FEBS Letters, 584(1), pp.199–202.

Potter, V.R. & Dubois, K.P., 1943. Studies on the Mechanism of Hydrogen Transport in Animal Tissues: VI. Inhibitor Studies with Succinic Dehydrogenase. The Journal of general physiology, 26(4), pp.391–404.

Preibisch, S., Saalfeld, S. & Tomancak, P., 2009. Globally optimal stitching of tiled 3D microscopic image acquisitions. Bioinformatics (Oxford, England), 25(11), pp.1463–1465.

Raymond, P.A. et al., 2014. Patterning the cone mosaic array in zebrafish retina requires specification of ultraviolet-sensitive cones. PLoS ONE, 9(1), p.e85325.

Raymond, P.A., Barthel, L.K. & Curran, G.A., 1995. Developmental patterning of rod and cone photoreceptors in embryonic zebrafish. The Journal of Comparative Neurology, 359(4), pp.537–550.

Rea, R. et al., 2004. Streamlined synaptic vesicle cycle in cone photoreceptor terminals. Neuron, 41(5), pp.755–766.

Reddy, A.B. et al., 2006. Circadian orchestration of the hepatic proteome. Current biology: CB, 16(11), pp.1107–1115.

Refinetti, R., Lissen, G.C. & Halberg, F., 2007. Procedures for numerical analysis of circadian rhythms. Biological rhythm research, 38(4), pp.275–325.

Rodríguez-Muela, N. et al., 2015. Lysosomal membrane permeabilization and autophagy blockade contribute to photoreceptor cell death in a mouse model of retinitis pigmentosa. Cell Death and Differentiation, 22(3), pp.476–487.

Sajdak, B.S. et al., 2019. Evaluating seasonal changes of cone photoreceptor structure in the 13-lined ground squirrel. Vision research, 158, pp.90–99.

Salvi, M. et al., 2007. Identification of the flavoprotein of succinate dehydrogenase and aconitase as in vitro mitochondrial substrates of Fgr tyrosine kinase. FEBS Letters, 581(29), pp.5579–5585.

Sartori-Rupp, A. et al., 2019. Correlative cryo-electron microscopy reveals the structure of TNTs in neuronal cells. Nature Communications, 10(1), p.342.

Scarpulla, R.C., Vega, R.B. & Kelly, D.P., 2012. Transcriptional integration of mitochondrial biogenesis. Trends in endocrinology and metabolism: TEM, 23(9), pp.459–466.

Simpson, C.F. & Kling, J.M., 1968. The mechanism of mitochondrial extrusion from phenylhydrazine-induced reticulocytes in the circulating blood. The Journal of cell biology, 36(1), pp.103–109.

Slautterback, D.B., 1965. Mitochondria in Cardiac Muscle Cells of the Canary and Some Other Birds. The Journal of cell biology, 24, pp.1–21.

Sloat, S. et al., 2016. Quantification of Mitochondrial Structure in Photoreceptors. Investigative Ophthalmology & Visual Science, 57(12).

Soubannier, V. et al., 2012. A vesicular transport pathway shuttles cargo from mitochondria to lysosomes. Current biology: CB, 22(2), pp.135–141.

Stone, J. et al., 2008. The locations of mitochondria in mammalian photoreceptors: Relation to retinal vasculature. Brain Research, 1189, pp.58–69.

Strappazzon, F. et al., 2015. AMBRA1 is able to induce mitophagy via LC3 binding, regardless of PARKIN and p62/SQSTM1. Cell Death and Differentiation, 22(3), pp.419–432.

Sugiura, A. et al., 2014. A new pathway for mitochondrial quality control: mitochondrial-derived vesicles. The EMBO Journal, 33(19), pp.2142–2156.

Sugiura, A. et al., 2017. Newly born peroxisomes are a hybrid of mitochondrial and ER-derived pre-peroxisomes. Nature, 542(7640), pp.251–254.

Tarboush, R. et al., 2014. Variability in mitochondria of zebrafish photoreceptor ellipsoids. Visual Neuroscience, 31(1), pp.11–23.

Tarlow, D.M. et al., 1977. Lipogenesis and the synthesis and secretion of very low density lipoprotein by avian liver cells in nonproliferating monolayer culture. Hormonal effects. The Journal of cell biology, 73(2), pp.332–353.

Tsukamoto, Y. et al., 2001. Microcircuits for night vision in mouse retina. Journal of Neuroscience, 21(21), pp.8616–8623.

Tyrrell, L.P. et al., 2019. A novel cellular structure in the retina of insectivorous birds. Scientific Reports, 9(1), p.15230.

Unuma, K. et al., 2015. Extrusion of mitochondrial contents from lipopolysaccharide-stimulated cells: Involvement of autophagy. Autophagy, 11(9), pp.1520–1536.

Urschel, M.R. & O’Brien, K.M., 2008. High mitochondrial densities in the hearts of Antarctic icefishes are maintained by an increase in mitochondrial size rather than mitochondrial biogenesis. Journal of Experimental Biology, 211(16), pp.2638–2646.

Utsumi, S. et al., 2020. Presence of ES1 homolog in the mitochondrial intermembrane space of porcine retinal cells. Biochemical and Biophysical Research Communications.

Van Laar, V.S. et al., 2018. Evidence for Compartmentalized Axonal Mitochondrial Biogenesis: Mitochondrial DNA Replication Increases in Distal Axons As an Early Response to Parkinson’s Disease-Relevant Stress. Journal of Neuroscience, 38(34), pp.7505–7515.

Vatine, G. et al., 2011. It’s time to swim! Zebrafish and the circadian clock. FEBS Letters, 585(10), pp.1485–1494.

Vincent, A.E. et al., 2019. Quantitative 3D Mapping of the Human Skeletal Muscle Mitochondrial Network. CellReports, 26(4), pp.996–1009.e4.

Wang, M. et al., 2015. The zebrafish period2 protein positively regulates the circadian clock through mediation of retinoic acid receptor (RAR)-related orphan receptor a (Rora). Journal of Biological Chemistry, 290(7), pp.4367–4382.

Wang, X. & Gerdes, H.-H., 2015. Transfer of mitochondria via tunneling nanotubes rescues apoptotic PC12 cells. Cell Death and Differentiation, 22(7), pp.1181–1191.

Wen, X., Saltzgaber, G.W. & Thoreson, W.B., 2017. Kiss-and-Run Is a Significant Contributor to Synaptic Exocytosis and Endocytosis in Photoreceptors. Frontiers in cellular neuroscience, 11, p.286.

Winkler, B.S., 1981. Glycolytic and oxidative metabolism in relation to retinal function. The Journal of general physiology, 77(6), pp.667–692.

Wittig, D. et al., 2012. Multi-level communication of human retinal pigment epithelial cells via tunneling nanotubes. PLoS ONE, 7(3), p.e33195.

Wong, Y.C., Peng, W. & Krainc, D., 2019. Lysosomal Regulation of Inter-mitochondrial Contact Fate and Motility in Charcot-Marie-Tooth Type 2. Developmental Cell, 50(3), pp.339–354.e4.

Youle, R.J. & Narendra, D.P., 2011. Mechanisms of mitophagy. Nature Reviews Molecular Cell Biology, 12(1), pp.9–14.

