## Supplemental Information for "Daily mitochondrial dynamics in cone photoreceptors"

**Author List Footnotes**
[1]Department of Biochemistry, University of Washington, Seattle, WA, USA
[2]Department of Ophthalmology, University of Washington, Seattle, WA, USA

**Supplemental Information**
- 5 Supplemental Figures
- 4 Tables
- 4 Videos
- Materials and methods

#### Supplemental Figures

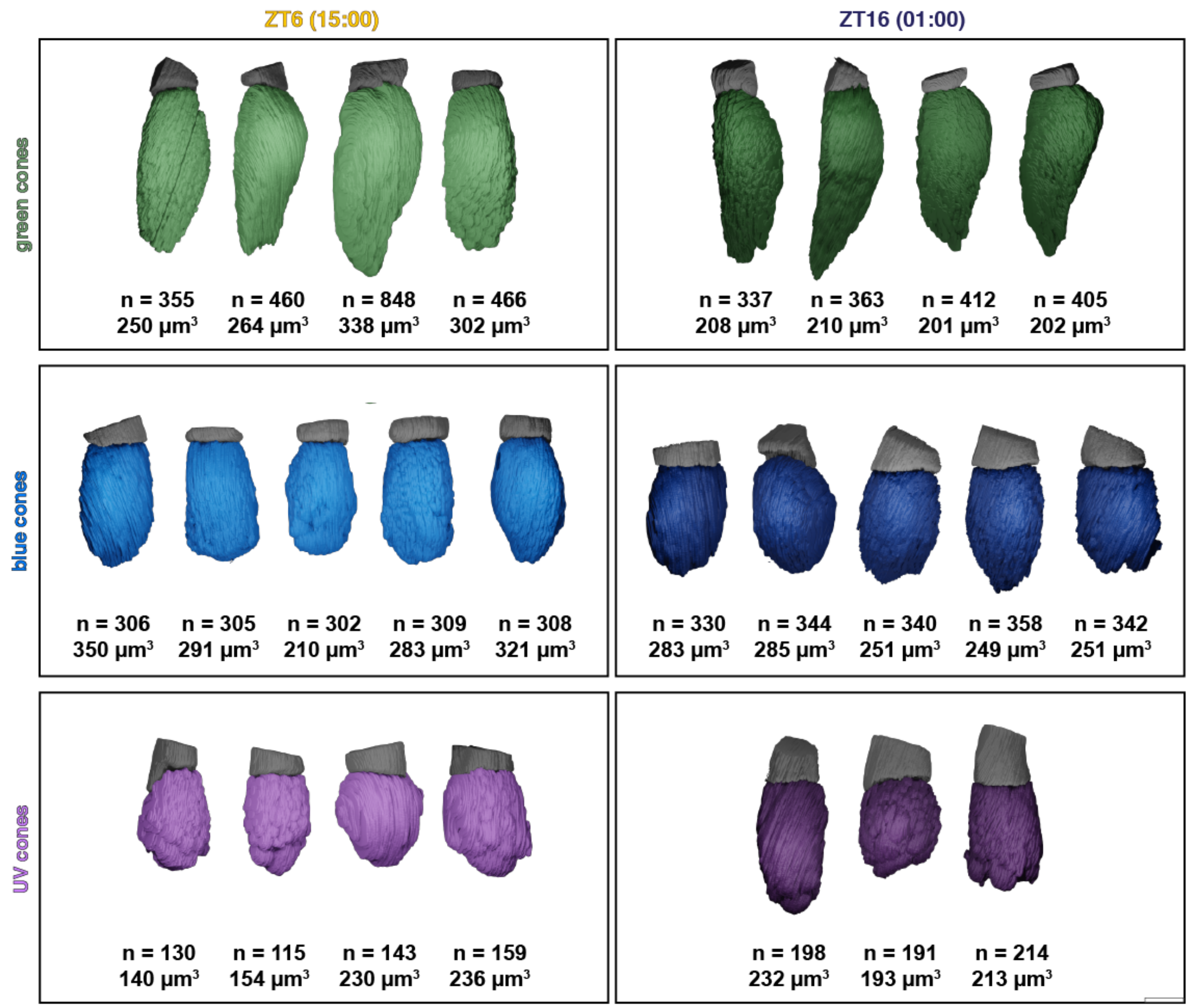


**Figure S1.** Gallery of 3-D renderings for all SBFSEM cells used in this study. Ellipsoids and OS are shown for green (top), blue (middle), and UV (bottom) cones, at 15:00LD (ZT6, left) and 01:00LD (ZT16, right). Corresponding mitochondrial numbers (n =) and total cluster volumes (in $\mu m^3$) are listed below each cell. Scale bar, 5 µm.

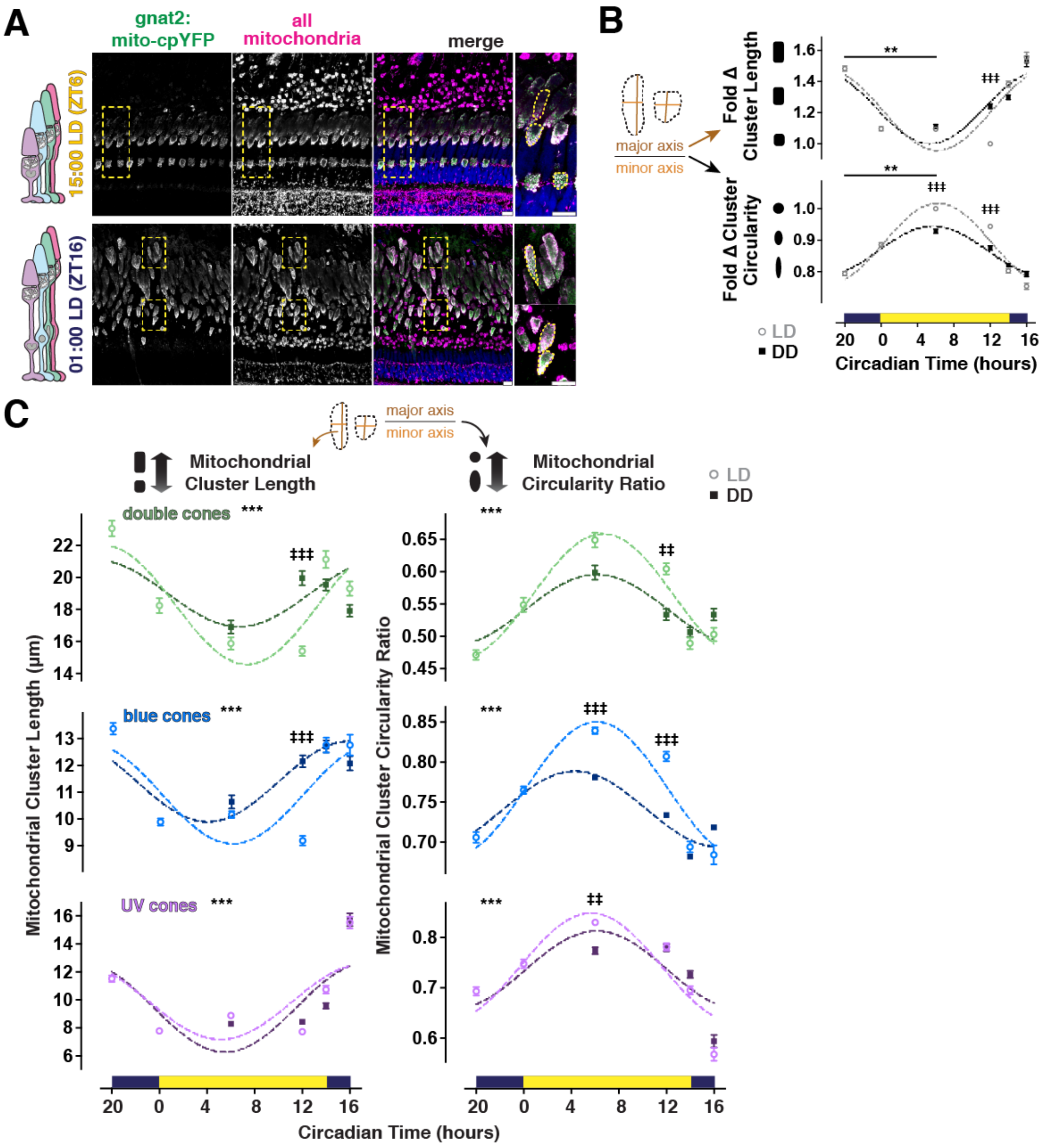


**Figure S2.** IHC morphological analysis of cone mitochondrial clusters throughout the day. A) IHC images of transgenic zebrafish outer retina expressing cone-targeted mito-cpYFP (green, left panels) overlaid with mitochondrial and nuclear stains (magenta and blue, respectively). Top row, light (ZT6); bottom row, dark (ZT16). Yellow boxes indicate zoomed-in area and tracings of single clusters; scale bars, 10 µm. B) Quantification of mean mitochondrial cluster length (left) and circularity ratio (right) from IHC over time. All cones at each timepoint were normalized and pooled for light-adapted (LD, grey open circles) or dark-adapted (DD, black squares) groups. C) Quantification of mean mitochondrial cluster length (left) and circularity ratio (right) from IHC over time for red-green double (top), blue (middle), and UV (bottom) cones in both LD or DD groups. Table 2 lists p-values and Ns from each group.

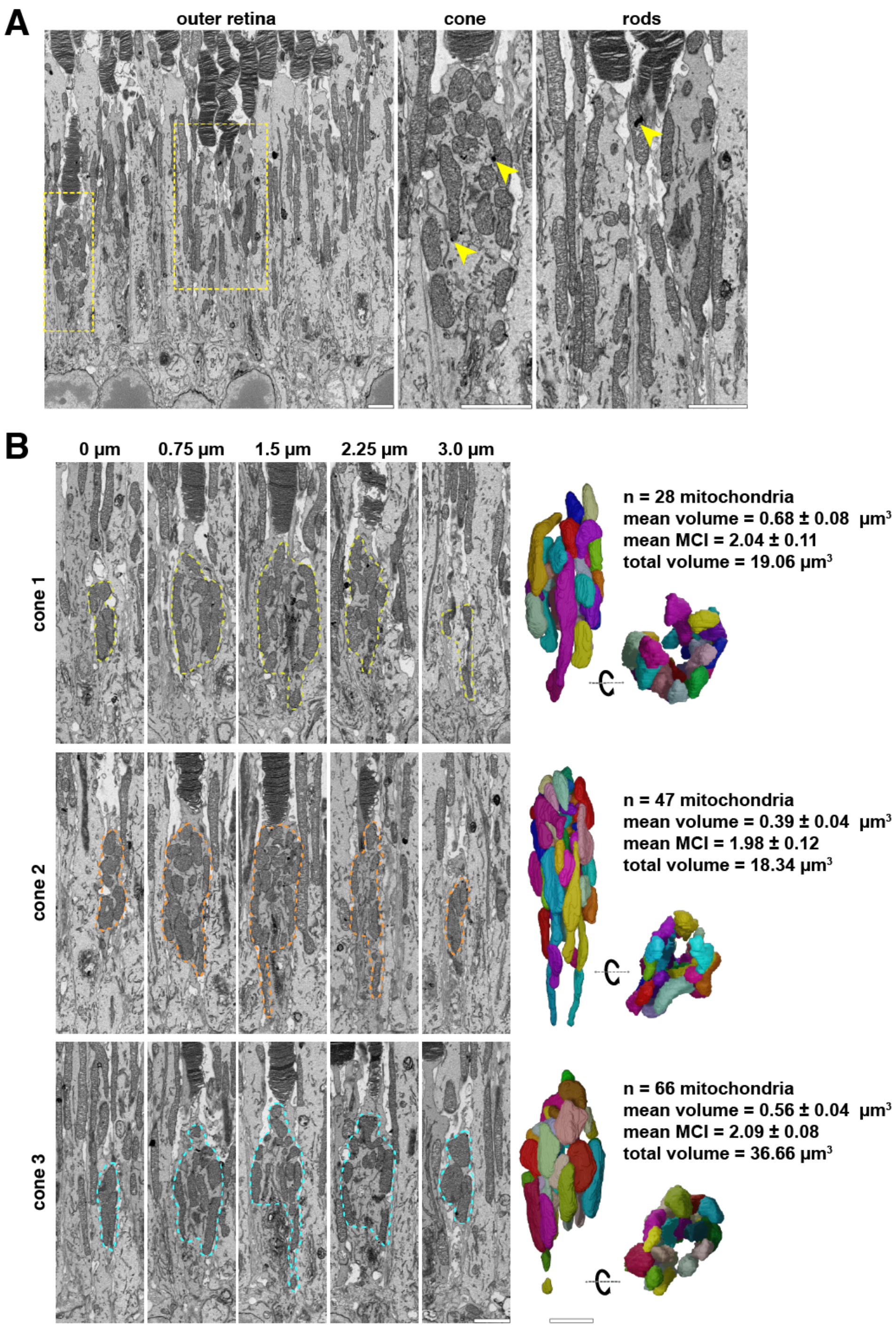


**Figure S3.** Mouse cone mitochondria have deposits and reside in heterogeneous clusters. A) SBFSEM images of C57BL/6J mouse outer retina; yellow boxes indicate zoomed-in areas. Cones (middle) and rods (right) display dark deposits (yellow arrowheads). B) Z-stacks from SBFSEM with 3-D rendered mitochondrial clusters (colored) from 3 individual mouse cones, and corresponding morphological measures ± S.E.M. Scale bars, 2 µm.

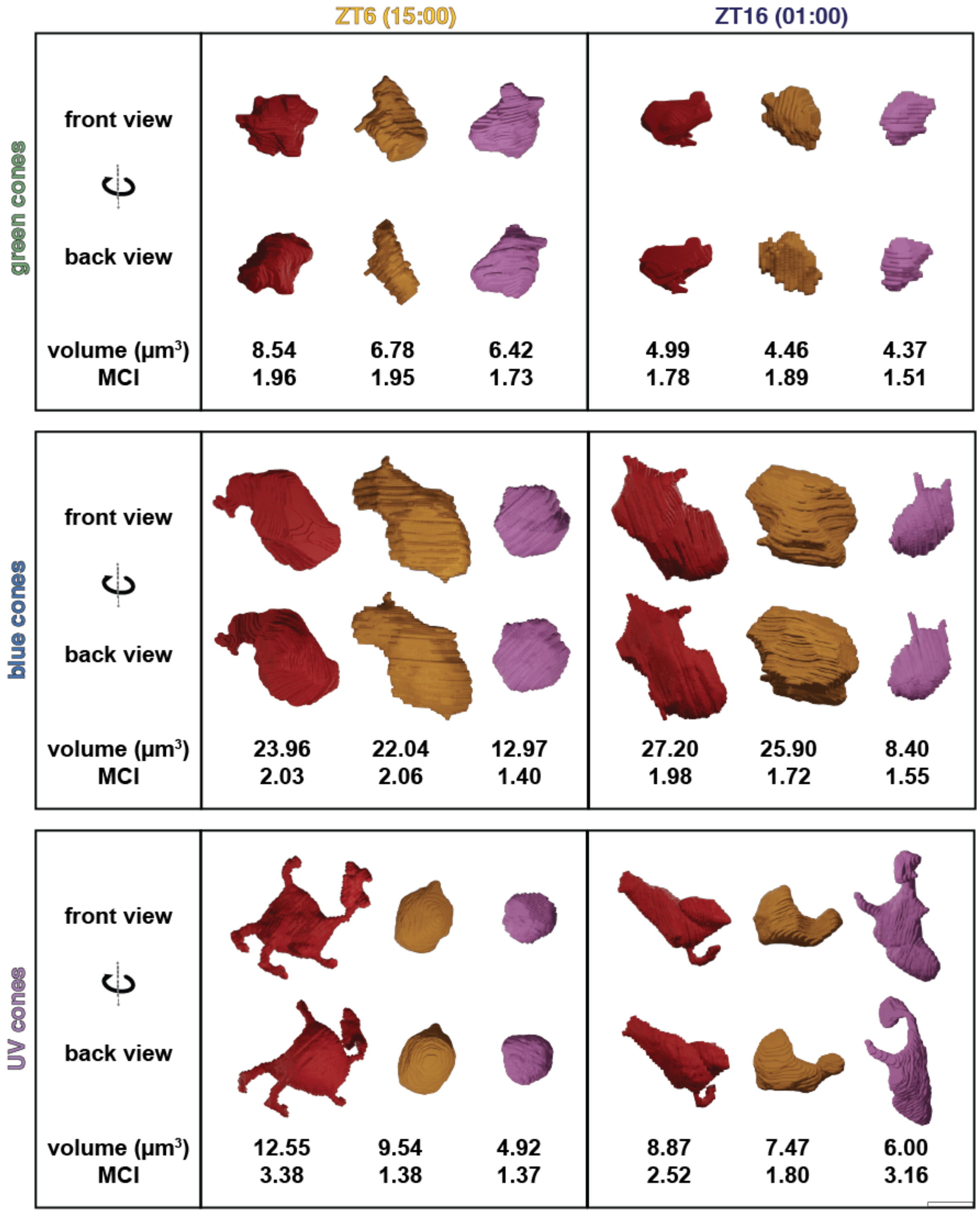


**Figure S4.** Gallery of megamitochondria in cone subtypes in day and night. Each mitochondrion from Figure 3A is represented in 3-D, viewed from opposite sides, with corresponding volume and MCI information. Green (top), blue (middle), and UV (bottom) cones are shown at ZT6 (left) and ZT16 (right). Scale bar, 2 μm.

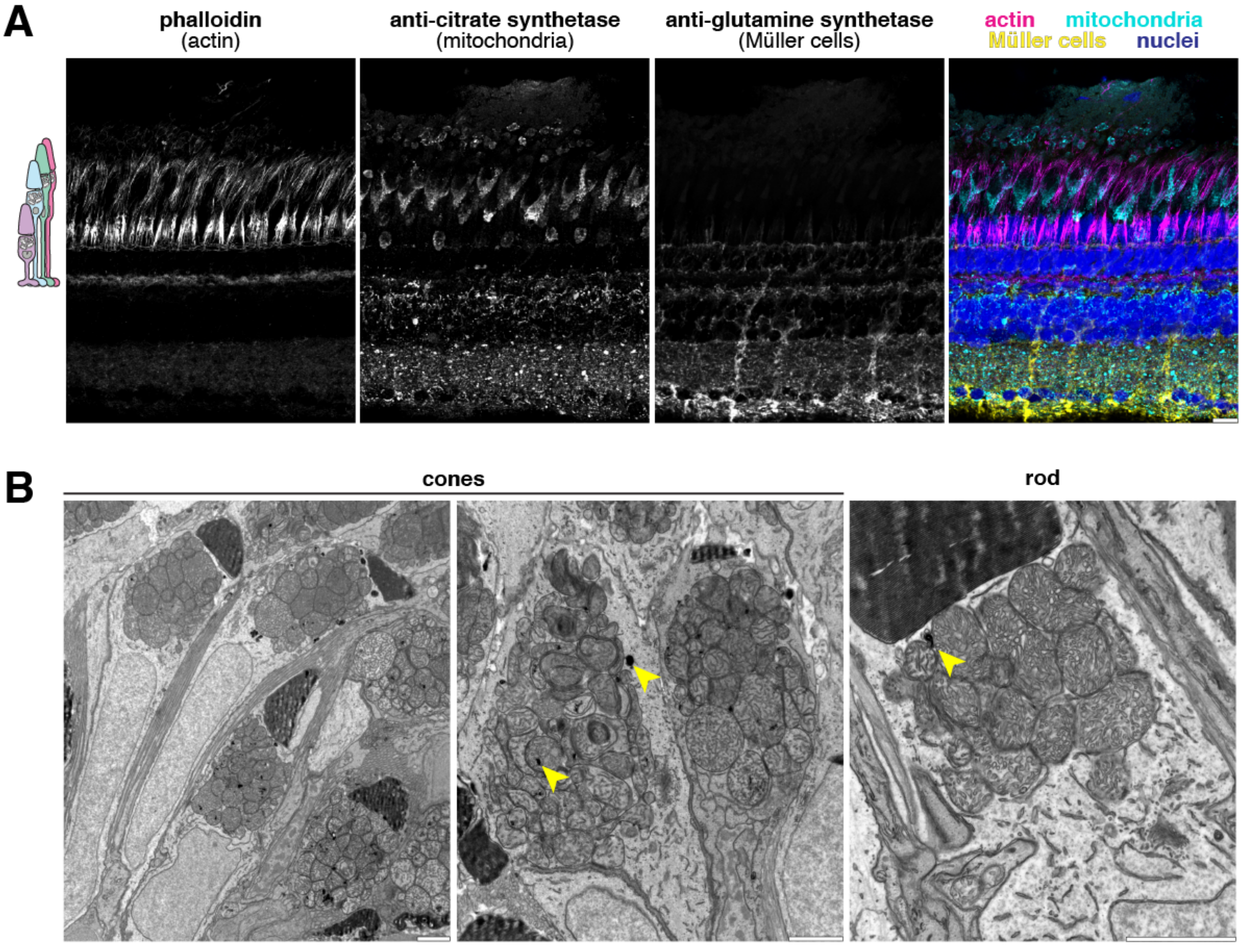


**Figure S5.** Albino zebrafish do not undergo retinal degeneration, and their cones have mitochondrial deposits. A) IHC images of light-adapted adult albino zebrafish retina. Phalloidin labels actin (magenta), anti-citrate synthetase labels all mitochondria (cyan), anti-glutamine synthetase labels Müller cells (yellow), and markers are counterstained with Hoechst to label nuclei (blue). Scale bar represents 10 μm. B) TEM images of albino zebrafish photoreceptors displaying normal morphology and dark deposits (yellow arrowheads). Scale bars, 2 μm.

### Tables

#### Table 1 – Antibodies and stains

**Antibodies and Stains**

| Target | Description | Species | Manufacturer | Catalog # | RRID # | Dilution |
|---|---|---|---|---|---|---|
| GFP | 1º Antibody | chicken | Abcam | ab13970 | RRID:AB_300798 | 1:10,000 |
| MTCO1 | 1º Antibody | mouse | Abcam | ab14705 | RRID:AB_2084810 | 1:1,000 |
| SDHB | 1º Antibody | mouse | Abcam | ab14714 | RRID:AB_301432 | 1:1,000 |
| ATP5A | 1º Antibody | mouse | Abcam | ab110273 | RRID:AB_10858175 | 1:1,000 |
| glutamine synthetase | 1º Antibody | mouse | Millipore | MAB302 | RRID:AB_2110656 | 1:1,000 |
| citrate synthetase | 1º Antibody | rabbit | Abcam | ab96600 | RRID:AB_10678258 | 1:500 |
| chicken IgY | 2º Antibody, AlexaFluor 488 conjugate | goat | Invitrogen | A11039 | RRID:AB_2534096 | 1:2,000 |
| mouse IgG | 2º Antibody, AlexaFluor 488 conjugate | goat | Invitrogen | A11001 | RRID:AB_2534069 | 1:2,000 |
| mouse IgG | 2º Antibody, AlexaFluor 647 conjugate | goat | Invitrogen | A21236 | RRID:AB_2535805 | 1:2,000 |
| rabbit IgG | 2º Antibody, AlexaFluor 647 conjugate | goat | Invitrogen | A32733 | RRID:AB_2633282 | 1:2,000 |
| cone outer segments | PNA stain, AlexaFluor 647 conjugate | | Invitrogen | L32460 | | 1:8,000 |
| actin | phalloidin stain, AlexaFluor 568 conjugate | | Invitrogen | A12380 | | 1:200 |
| nuclei | Hoechst stain | | Invitrogen | H3569 | | 10 μM |

### Table 2 – Experimental sample numbers by figure

**Experimental Sample Numbers and Statistics by Figure**

| figure | description | cell type | timepoint | # animals | # technical replicates | # cells | # mito-chondria | error bars = | statistical test(s) | p-values for * | p-values for ‡ |
|---|---|---|---|---|---|---|---|---|---|---|---|
| 1D | SBFSEM dot quantification | green | 1500LD | 2 | 2 | 4 | | | Brown-Forsythe ANOVA, Welch's unpaired t test | Between cone types at ZT6, * p<0.05, ** p<0.01, *** p<0.001 | Between ZT6 and ZT16, ‡ p<0.05, ‡‡ p<0.01, ‡‡‡ p<0.001 |
| | | | 0100LD | 2 | 2 | 4 | | | | | |
| | | blue | 1500LD | 2 | 2 | 5 | | | | | |
| | | | 0100LD | 2 | 2 | 5 | | | | | |
| | | UV | 1500LD | 2 | 2 | 4 | | | | | |
| | | | 0100LD | 2 | 2 | 3 | | | | | |
| 2B | MCI v. volume | all | light | 2 | 2 | 4 | 1187 | | | | |
| | | | dark | 2 | 2 | 5 | 1692 | | | | |
| 2E, 2F | MCI, volume histograms from manual segmentation | green | 1500LD | 1 | 1 | 1 | 464 | | Kolmogorov-Smirnov test | | |
| | | | 0100LD | 2 | 2 | 2 | 827 | | | | |
| | | blue | 1500LD | 2 | 2 | 2 | 612 | | | | |
| | | | 0100LD | 2 | 2 | 2 | 675 | | | | |
| | | UV | 1500LD | 1 | 1 | 1 | 115 | | | | |
| | | | 0100LD | 1 | 1 | 1 | 190 | | | | |
| 3C, 3D | point sizes, core-periphery and bottom-top | green | 1500LD | 2 | 2 | 4 | 2101 | SD | Ordinary one-way ANOVA, Mann-Whitney unpaired t test | For regional comparisons, * p<0.05, ** p<0.01, *** p<0.0001 | Between ZT6 and ZT16, ‡ p<0.05, ‡‡ p<0.001, ‡‡‡ p<0.0001 |
| | | | 0100LD | 2 | 2 | 4 | 1223 | | | | |
| | | blue | 1500LD | 2 | 2 | 5 | 1528 | | | | |
| | | | 0100LD | 2 | 2 | 5 | 1714 | | | | |
| | | UV | 1500LD | 2 | 2 | 4 | 543 | | | | |
| | | | 0100LD | 2 | 2 | 3 | 590 | | | | |
| 3F | er-GFP IHC | blue and UV | 500 | 2 | 2 | 110 | | SEM | Kruskal-Wallis test with Dunn's multiple comparisons | For timepoint comparisons to ZT0 in LD group, ** p<0.01, *** p<0.0001 | Between LD and DD groups, ‡ p<0.05, ‡‡ p<0.001, ‡‡‡ p<0.0001 |
| | | | 900 | 2 | 2 | 101 | | | | | |
| | | | 1500LD | 2 | 2 | 127 | | | | | |
| | | | 1500DD | 2 | 2 | 110 | | | | | |
| | | | 2100LD | 2 | 2 | 101 | | | | | |
| | | | 2100DD | 2 | 2 | 115 | | | | | |
| | | | 2300LD | 2 | 2 | 129 | | | | | |
| | | | 2300DD | 2 | 2 | 86 | | | | | |
| | | | 0100LD | 2 | 2 | 97 | | | | | |
| | | | 0100DD | 2 | 2 | 119 | | | | | |
| 4B, 4C | qPCR | retina | all | 3 | 3 | | | SEM | 2-way ANOVA with Tukey's multiple comparisons | For peak-trough comparisons, * p<0.05, ** p<0.01, *** p<0.0001 | |
| 4E | LC3-GFP IHC | blue and UV | 500 | 2 | 2 | 145 | | SEM | Ordinary one-way ANOVA with Dunnett's multiple comparisons | For timepoint comparisons to ZT20 in LD group, *** p<0.0001 | Between LD and DD groups, ‡‡ p<0.001, ‡‡‡ p<0.0001 |
| | | | 900 | 2 | 2 | 152 | | | | | |
| | | | 1500LD | 2 | 2 | 160 | | | | | |
| | | | 1500DD | 2 | 2 | 77 | | | | | |
| | | | 2100LD | 2 | 2 | 159 | | | | | |
| | | | 2100DD | 2 | 2 | 144 | | | | | |
| | | | 2300LD | 2 | 2 | 157 | | | | | |
| | | | 2300DD | 2 | 2 | 142 | | | | | |
| | | | 0100LD | 2 | 2 | 146 | | | | | |
| | | | 0100DD | 2 | 2 | 152 | | | | | |
| 4F | SBFSEM lucent structures | all | light | 2 | 2 | 24 | | | Mann-Whitney test | | For light-dark comparison, ‡‡‡ p<0.0001 |
| | | | dark | 2 | 2 | 18 | | | | | |
| 5E | deposits, exocytosis | all | light | 2 | 2 | 24 | | | Mann-Whitney test | | |
| | | | dark | 2 | 2 | 18 | | | | | |
| 6C | SDH histochemistry | all | 500 | 3 | 2 | 104 | | SEM | Kruskal-Wallis test with Dunn's multiple comparisons | Between ZT0 and ZT14 in LD group, ** p<0.001 | Between LD and DD groups, ‡ p<0.05, ‡‡ p<0.01 |
| | | | 900 | 3 | 2 | 90 | | | | | |
| | | | 1500LD | 3 | 2 | 77 | | | | | |
| | | | 1500DD | 3 | 2 | 116 | | | | | |
| | | | 2100LD | 3 | 2 | 65 | | | | | |
| | | | 2100DD | 3 | 2 | 113 | | | | | |
| | | | 2300LD | 3 | 2 | 119 | | | | | |
| | | | 2300DD | 3 | 2 | 57 | | | | | |
| | | | 0100LD | 3 | 2 | 50 | | | | | |
| | | | 0100DD | 3 | 2 | 89 | | | | | |
| 6C | COX histochemistry | all | 500 | 3 | 2 | 95 | | | | | |
| | | | 900 | 3 | 2 | 95 | | | | | |
| | | | 1500LD | 3 | 2 | 80 | | | | | |
| | | | 1500DD | 3 | 2 | 105 | | | | | |
| | | | 2100LD | 3 | 2 | 79 | | | | | |
| | | | 2100DD | 3 | 2 | 105 | | | | | |
| | | | 2300LD | 3 | 2 | 119 | | | | | |
| | | | 2300DD | 3 | 2 | 113 | | | | | |
| | | | 0100LD | 3 | 2 | 115 | | | | | |
| | | | 0100DD | 3 | 2 | 120 | | | | | |
| 6D, 6E | ${}^{13}$C succinate, light | retina | 0m | 3 | 3 | | | SD | 2-way ANOVA with Tukey's multiple comparisons | | Between light-dark groups, ‡ p<0.05, ‡‡ p<0.001, ‡‡‡ p<0.0001 |
| | | | 1m | 3 | 3 | | | | | | |
| | | | 3m | 3 | 3 | | | | | | |
| | | | 5m | 5 | 5 | | | | | | |
| | | | 10m | 3 | 3 | | | | | | |
| | | | 60m | 3 | 3 | | | | | | |
| | ${}^{13}$C succinate, dark | retina | 0m | 3 | 3 | | | | | | |
| | | | 1m | 3 | 3 | | | | | | |
| | | | 3m | 3 | 3 | | | | | | |
| | | | 5m | 5 | 5 | | | | | | |
| | | | 10m | 3 | 3 | | | | | | |
| | | | 60m | 3 | 3 | | | | | | |
| 6F | ${}^{13}$C succinate, light | media | 60m | 3 | 3 | | | SD | 2-way ANOVA with Sidak's multiple comparisons | | |
| | ${}^{13}$C succinate, dark | media | 60m | 3 | 3 | | | | | | |
| S2B | mito-cpYFP IHC | all | 500 | 2 | 3 | 317 | | SEM | Kruskal-Wallis test with Dunn's multiple comparisons | For peak-trough comparisons in the LD group, ** p<0.001, *** p<0.0001 | Between LD and DD groups, ‡‡ p<0.001, ‡‡‡ p<0.0001 |
| | | | 900 | 2 | 3 | 321 | | | | | |
| | | | 1500LD | 2 | 3 | 341 | | | | | |
| | | | 1500DD | 2 | 3 | 226 | | | | | |
| | | | 2100LD | 2 | 3 | 325 | | | | | |
| | | | 2100DD | 2 | 3 | 294 | | | | | |
| | | | 2300LD | 2 | 3 | 284 | | | | | |
| | | | 2300DD | 2 | 3 | 348 | | | | | |
| | | | 0100LD | 2 | 3 | 256 | | | | | |
| | | | 0100DD | 2 | 3 | 276 | | | | | |
| S2C | mito-cpYFP IHC | double | 500 | 2 | 3 | 124 | | | | | |
| | | | 900 | 2 | 3 | 114 | | | | | |
| | | | 1500LD | 2 | 3 | 117 | | | | | |
| | | | 1500DD | 2 | 3 | 72 | | | | | |
| | | | 2100LD | 2 | 3 | 138 | | | | | |
| | | | 2100DD | 2 | 3 | 111 | | | | | |
| | | | 2300LD | 2 | 3 | 106 | | | | | |
| | | | 2300DD | 2 | 3 | 123 | | | | | |
| | | | 0100LD | 2 | 3 | 98 | | | | | |
| | | | 0100DD | 2 | 3 | 94 | | | | | |
| | mito-cpYFP IHC | blue | 500 | 2 | 3 | 94 | | | | | |
| | | | 900 | 2 | 3 | 120 | | | | | |
| | | | 1500LD | 2 | 3 | 116 | | | | | |
| | | | 1500DD | 2 | 3 | 78 | | | | | |
| | | | 2100LD | 2 | 3 | 107 | | | | | |
| | | | 2100DD | 2 | 3 | 91 | | | | | |
| | | | 2300LD | 2 | 3 | 96 | | | | | |
| | | | 2300DD | 2 | 3 | 117 | | | | | |
| | | | 0100LD | 2 | 3 | 68 | | | | | |
| | | | 0100DD | 2 | 3 | 80 | | | | | |
| | mito-cpYFP IHC | UV | 500 | 2 | 3 | 99 | | | | | |
| | | | 900 | 2 | 3 | 87 | | | | | |
| | | | 1500LD | 2 | 3 | 108 | | | | | |
| | | | 1500DD | 2 | 3 | 76 | | | | | |
| | | | 2100LD | 2 | 3 | 80 | | | | | |
| | | | 2100DD | 2 | 3 | 92 | | | | | |
| | | | 2300LD | 2 | 3 | 82 | | | | | |
| | | | 2300DD | 2 | 3 | 108 | | | | | |
| | | | 0100LD | 2 | 3 | 90 | | | | | |
| | | | 0100DD | 2 | 3 | 102 | | | | | |

#### Table 3 – Primers used for qPCR

**Primer Sequences Used for qPCR**

| Gene | Forward Primer | Reverse Primer |
|---|---|---|
| *pgc1α* | 5'-GCCGTTCTCCCTCTTGGTCT-3' | 5'-AGACGCTCGTGCTGGTATTC-3' |
| *pgc1β* | 5'-CTCCTGTTGACACCCCCTAA-3' | 5'-TGGAGTTCAGTGCAAAGACG-3' |
| *nrf1* | 5'-CAGTCGGAGCGTTAACTGGA-3' | 5'-ATTCGAGTCGTTACGGGAGC-3' |
| *tfam* | 5'-GGGAACTACTACGCAGGCAA-3' | 5'-GTGCTCCTCCCACGATTTGA-3' |
| *polg1* | 5'-GGGAAACTTTCAGGAGCTGATGC-3' | 5'-CCAGCATACCAGCAAGAGTCA-3' |
| *sirt3* | 5'-TGCTCAGGATGTACACGCAAA-3' | 5'-TGTCATCCCGAAGTTCCTCTCC-3' |
| *mfn2* | 5'-TGCTCCAGTGCAATAACCGT-3' | 5'-TCTTATCCACCTGCTCCCGA-3' |
| *es1* | 5'-TCACAACACCCGTGAACCTT-3' | 5'-GGTCTCCCACATGAAGGTCG-3' |
| *aanat2* | 5'-AGAACAGGAGGCTATGAGCAC-3' | 5'-CGCAGGTACTGGAGGTAACG-3' |

#### Table 4 – Fitting parameters for cosinor curves

**Fitting Parameters for Cosinor Curves** $Y(t) = M + A\cos(\Omega + \Phi)$

| Curve | Condition | MESOR (M) | Amplitude (A) | Phaseshift (Φ) | Fit ($R^2$) | Max Fold Δ |
|---|---|---|---|---|---|---|
| IHC - ER-Mitochondrial Apposition Ratio | LD | 0.682 | 0.173 | 9.82 | 0.258 | 1.67 |
| | DD | 0.612 | 0.107 | 10.21 | 0.129 | 1.62 |
| qPCR - *pgc1α* | LD | 1.044 | 0.384 | 4.92 | 0.340 | 2.91 |
| | DD | 1.221 | 0.244 | 10.69 | 0.129 | 2.47 |
| qPCR - *pgc1β* | LD | 1.044 | 0.195 | 10.96 | 0.246 | 1.45 |
| | DD | 1.130 | 0.278 | 10.83 | 0.306 | 2.30 |
| qPCR - *nrf1* | LD | 1.294 | 0.345 | 20.06 | 0.448 | 2.10 |
| | DD | 1.427 | 0.518 | 20.04 | 0.452 | 2.61 |
| qPCR - *tfam* | LD | 1.606 | 0.557 | -16.96 | 0.261 | 2.87 |
| | DD | 2.021 | 0.820 | 14.68 | 0.258 | 3.89 |
| qPCR - *polg1* | LD | 3.445 | 2.808 | 15.02 | 0.554 | 7.04 |
| | DD | 3.815 | -3.361 | 18.26 | 0.512 | 7.41 |
| qPCR - *sirt3* | LD | 1.772 | -0.477 | 18.24 | 0.213 | 2.36 |
| | DD | 2.213 | 0.849 | 15.08 | 0.448 | 2.93 |
| qPCR - *mfn2* | LD | 0.864 | 0.432 | -0.82 | 0.483 | 4.83 |
| | DD | 0.853 | 0.417 | -0.70 | 0.418 | 3.72 |
| qPCR - *es1* | LD | 1.390 | 0.636 | 15.97 | 0.417 | 2.06 |
| | DD | 1.526 | 0.664 | 15.70 | 0.126 | 2.47 |
| qPCR - *aanat2* | LD | 4.357 | -4.217 | 17.06 | 0.722 | 89.81 |
| | DD | 6.273 | -6.721 | -1.67 | 0.678 | 112.49 |
| IHC - Mitochondrial-Associated Autophagosomes | LD | 2.222 | 1.060 | 10.33 | 0.166 | 2.99 |
| | DD | 2.632 | 0.828 | 9.83 | 0.066 | 1.57 |
| Histochemistry - SDH activity | LD | 1.108 | 0.180 | 13.44 | 0.022 | 1.30 |
| | DD | 1.176 | 0.167 | 15.17 | 0.004 | 1.12 |
| Histochemistry - COX activity | LD | 0.970 | -0.007 | 2.97 | 0.001 | 1.06 |
| | DD | 1.015 | 0.040 | 11.45 | 0.010 | 1.03 |

#### Videos

**Video 1** – Example 3-D SBFSEM stack of single cone mitochondrial cluster at 100 nm Z-resolution. The same cell is shown as a raw image, and analyzed by rapid dot quantification or manual segmentation for the corresponding 3-D renderings. Scale bar, 2 μm.

**Video 2** – 3-D renderings of cone mitochondrial clusters analyzed by manual segmentation. Shown are clusters from green (left), blue (middle) and UV (right) cones at 15:00 (ZT6, top) and 01:00 (ZT16, bottom). Scale bar, 5 μm.

**Video 3** – At night (ZT16) extruded mitochondrial material forms stalks and sacs in the extracellular space. Part 1) SBFSEM stack of a single cone mitochondrial cluster at 50 nm Z-resolution shown as a raw image, and analyzed by manual segmentation for the corresponding 3-D rendering. Yellow, plasma membrane; green, outer segment; reds, mitochondria; blues, mitochondrial deposits. Scale bar, 2 μm. Part 2) 3-D rendering of the structure in (1). Beige, plasma membrane; grey, outer segment; reds, mitochondria; blues, mitochondrial deposits.

**Video 4** – At night (ZT16) extruded mitochondrial material forms extracellular networks around the cell surface. Part 1) SBFSEM stack of a single cone mitochondrial cluster at 50 nm Z-resolution shown as a raw image, and analyzed by manual segmentation for the corresponding 3-D rendering. Yellow, plasma membrane; green, outer segment; reds, mitochondria; blues, mitochondrial deposits. Scale bar, 2 μm. Part 2) 3-D rendering of the structure in (1). Beige, plasma membrane; grey, outer segment; reds, mitochondria; blues, mitochondrial deposits.

#### Materials and Methods

**Zebrafish maintenance and retina collection.** Research was authorized by the University of Washington Institutional Animal Care and Use Committee. Wild-type AB, transgenic *Roy*$^{-/-}$ and albino zebrafish were maintained in the University of Washington ISCRM aquatics facility at 27.5°C on a 14-hour/10-hour light/dark cycle. Fish used for experiments were male and female siblings between 6 and 12 months old. 24 h prior to experiments, fish were fasted in either continuous darkness (DD) or standard room light (LD). Fish designated for dark conditions or time points were euthanized and dissections were performed under infrared light. For qPCR and metabolic analyses using isolated retinas, fish from the daytime LD group were briefly dark-adapted 20-30 mins and eyes were dissected under red light to facilitate removal of the RPE. Fish were euthanized in an ice bath followed by cervical dislocation and enucleation.

**Generation of transgenic and albino zebrafish.** The transgenic lines Tg(gnat2:er-GFP) (George et al. 2014; RRID:ZDB-GENO-140416-8), Tg(gnat2:GFP-LC3) (George et al. 2014; RRID:ZDB-GENO-140416-6), and Tg(gnat2:mito-cpYFP) (Giarmarco et al. 2017; RRID:ZDB-GENO-181217-2) have been described previously. Transgenic lines were generated using the Gateway-Tol2 system (Kwan et al. 2007) to express fluorescent proteins downstream of the zebrafish cone transducin promoter *gnat2*. Transgenic fish used in this study were heterozygotes outcrossed from stable transgenic lines. Albino fish arose from a spontaneous recessive mutation in the *Roy*$^{-/-}$ background, and lack all pigmentation similar to the *crystal* strain described in (Antinucci & Hindges 2016).

**Serial block-face scanning electron microscopy.** Wild-type AB or albino zebrafish eyes were enucleated and the anterior half was immediately dissected away into fixative (4% glutaraldehyde in 0.1 M sodium cacodylate buffer, pH 7.2). Tissue was fixed overnight at room temperature (RT), then stored overnight at 4°C. Samples were washed 4 times in sodium cacodylate buffer, post-fixed in osmium ferrocyanide (2% osmium tetroxide/3% potassium ferrocyanide in buffer) for 1 h on ice, washed, incubated in 1% thiocarbohydrazide for 20 min, and washed again. After incubation in 2% osmium tetroxide for 30 min at RT, samples were washed and en bloc stained with 1% aqueous uranyl acetate overnight at 4°C. Samples were finally washed and en bloc stained with Walton's lead aspartate for 30 min at 60°C, dehydrated in a graded ethanol series, and embedded in Durcupan resin. Unless stated otherwise, five washes with water were used for all wash steps. Resin blocks were mounted in a Zeiss Sigma VP or Zeiss Merlin Gemini 2 scanning electron microscopes fitted with a Gatan 3View2XP ultramicrotome apparatus. Serial sections were cut at 50 nm thickness and imaged with 5 nm pixel size in the outer retina, 100-200 µm from the optic nerve. Z-stack images of individual mitochondrial clusters were aligned, stitched (Preibisch et al. 2009), and measured using the TrakEM2 plugin for ImageJ (RRID:SCR_008954). SBFSEM analysis of C57BL/6J mouse cones was conducted using a previously acquired dataset (Kanow et al. 2017).

**Quantification of mitochondrial number, volume and complexity.** Mitochondrial clusters were analyzed using dot quantification or manual segmentation; mitochondria were defined as structures in the cluster with a double outer membrane and cristae. Additionally, total volume was determined for each cluster via manual segmentation, connecting outer mitochondrial membranes. For dot quantification, a uniform-sized dot was placed in the center of each mitochondrion throughout the Z-stack; average mitochondrial volume was calculated by dividing the number of mitochondria by the cluster volume. For manual segmentation, individual mitochondria were traced throughout the Z-stack and reconstructed in 3-D. Mitochondrial Complexity Index (MCI) was assessed using the following equation (Vincent et al. 2019):

$$\mathrm{MCI} = \frac{SA^3}{16\pi^2 V^2}$$

Where SA equals surface area and V equals volume. Mitochondrial volumes and MCIs were compared using a frequency distribution.

**Quantification of mitochondrial size across clusters.** Mitochondrial point size was approximated from dot quantification; larger or more branched mitochondria were represented by more dots and thus larger point sizes. For each cell, X-Y-Z coordinates from the center of every mitochondrion and corresponding point sizes were loaded into Igor Pro (RRID:SCR_000325). Mitochondrial positions were binned in relation to an X-Y-Z axis along the cluster center using the distance formula. The cluster core was defined as the inner 2/3 of the cluster, and the periphery as the outer 1/3. Similarly, top and bottom halves of the cluster were defined by distance to the OS and nucleus, respectively. Mitochondrial point size was then assayed relative to cluster position, and core/periphery and bottom/top ratios were used as measures of size distribution across clusters.

**Immunohistochemistry.** Albino or transgenic *Roy*$^{-/-}$ zebrafish eyes were placed into fixative (4% paraformaldehyde in PBS, pH 7.3), pierced across the cornea with a needle, and the vitreous cavity was flushed with fixative. Following overnight immersion at 4°C, *Roy*$^{-/-}$ eyes were rinsed in PBS, then bleached with 10% $H_2O_2$ in PBS overnight at 37°C to remove pigmentation of the RPE (this step was omitted for albino eyes). Eyes were then cryoprotected in sucrose, the anterior halves dissected away, and eyecups were embedded and frozen in OCT cryomolds. Eyecups were cryosectioned at 20 μm onto slides that contained one section from each timepoint for parallel staining and analysis. Sections were washed in PBS, then blocked for 30 min in blocking buffer (2% normal donkey serum, 2 mg/mL bovine serum albumin and 0.3% Triton X-100 in PBS). Primary antibodies diluted in blocking buffer were then applied to cryosections overnight at 4°C. Antibodies and stains are reported in Table 1. Secondary antibodies conjugated to AlexaFluor dyes were diluted at 1:2,000 in blocking buffer, and incubated on sections for 1 hr in darkness at RT. Phalloidin staining was conducted prior to antibody incubations for 30 min at RT using 1:200 phalloidin in 2% Triton X-100 in PBS; peanut agglutinin was diluted 1:8,000 in blocking buffer and incubated overnight with primary antibodies. All sections were counterstained with Hoechst 33342 (Invitrogen) in PBS, and mounted with SouthernBiotech Fluoromount-G (Fisher Scientific) under glass coverslips.

**IHC imaging and analysis.** Fluorescence of the outer retina in a region 100-200 µm from the optic nerve was visualized using a Leica SP8 confocal microscope with a 63X oil objective. Z-stack images were acquired every 0.3 µm at 2048 x 2048 pixel resolution and 12-bit depth using Leica LAS-X software (RRID:SCR_013673). Z-stacks from each timepoint were blindly analyzed using ImageJ software (RRID:SCR_002285) to quantify features of individual cells. Cell types were identified by nuclear morphology and position in the outer retina; Table 2 lists Ns from all groups. Representative images were post-processed with Leica Lightning deconvolution, and brightness and contrast were adjusted equally to ease visualization.
For mito-cpYFP retinas, the perimeter of the mitochondrial cluster was traced and measured using one slice from the center of the mitochondrial cluster. Circularity of the ellipsoid region was calculated using the following equation:

$$\text{Circularity Ratio} = \frac{Minor\ Axis}{Major\ Axis}$$

For er-GFP, Z-stacks were maximum intensity projected over 5 µm depth, and the length of ER visible from the base to the top of the counterstained mitochondrial cluster was traced and measured. ER:Mitochondrial apposition ratio was calculated for blue and UV cones using the following equation:

$$\text{Apposition Ratio} = \frac{ER\ Opposition\ Length}{Ellipsoid\ Major\ Axis}$$

For GFP-LC3, Z-stacks were processed to apply Gaussian blur (radius 3) to the GFP channel and then maximum intensity projected over 5 µm depth. GFP-positive puncta overlapping with the counterstained mitochondrial cluster were counted for blue and UV cones.

**Quantitative PCR.** Wild-type AB zebrafish eyes were dissected in PBS, and the retina and RPE were mechanically dissociated. Retinas were snap frozen in liquid $N_2$ and stored at -80°C. mRNA was extracted using a RNeasy Micro Kit (Qiagen). mRNA purity and concentration were determined using a ND-2000C microvolume spectrophotometer (ThermoFisher). 0.5-1 µg of mRNA was reverse transcribed into cDNA using iScriptTM Reverse Transcriptase Supermix (Bio- Rad). Negative controls contained no reverse transcriptase, and cDNA samples for each timepoint were stored at -20°C. Normfinder was used to calculate reference gene stability (TATA-box-binding factor). Primer sequences are reported in Table 3. cDNA was diluted 1:40 and qPCR was run for each timepoint in a 7500 Fast Real-Time PCR System (ThermoFisher). All other timepoints were compared relative to ZT0 using the Livak ΔΔCt method. The 7500 Fast Real-Time PCR Software was used to rule out statistical outliers for technical triplicates. Biological outliers were ruled out using Grub's test. Lastly a cosinor curve of rhythm was fit to the data points, using the following equation (Refinetti et al. 2007):

$$\mathrm{Y(t)} = M + Acos(\Omega + \Phi)$$

Where M equals MESOR (Midline-Estimating Statistic of Rhythm), A equals Amplitude, Ω equals $\frac{2\pi}{24h}(0.2618)$, and Φ equals Phaseshift. Table 4 lists fitting parameters for all cosinor curves; in graphs solid lines depicted fits of $R^2$>0.5 and dashed lines, $R^2$<0.5.

**Analysis of mitochondrial deposits and extrusion.** Mitochondrial deposits and associated structures were manually segmented using image stacks obtained with SBFSEM. Deposits were defined as dark lamellar material inside of mitochondria or the cluster; extracellular stalks connected sacs were identified by a distinct dark, hollow staining pattern. Mitochondrial deposits and extrusion events were blindly quantified for each cell by a ranking system. For deposits: (1) no mitochondrial deposits, (2) few mitochondrial deposits, (3) many mitochondrial deposits, (4) every mitochondrion having a deposit. For the number of extrusion events: (1) no events, (2) one event, (3) two or three events, (4) more than three events. Rankings were compared using a frequency distribution.

**Enzyme histochemistry.** Histochemical enzyme activity was assayed similar to previous studies in human retina (Andrews et al. 1999) and mouse retina (Chinchore et al. 2017). All assay reagents were acquired from Sigma-Aldrich unless noted. Albino zebrafish eyes were dissected in PBS to remove the lens, and eyecups were immediately embedded and frozen in OCT cryomolds. Eyecups were cryosectioned at 14 µm onto frozen slides that contained one unfixed section from each time point for parallel staining and analysis. The same day, slides were stained for 10 min at 37°C. To assay SDH activity, sections were incubated in SDH reaction mixture (13 mM sodium succinate, 1.5 mM nitro blue tetrazolium, 0.2 mM phenazine methosulphate, and 1.0 mM sodium azide in PBS, pH 7.0) or negative control mixture (SDH reaction mixture lacking sodium succinate). To assay COX activity, sections were incubated in COX reaction mixture (4 mM 3 ,3'-diaminobenzidine hydrochloride, 6 mg/mL reduced cytochrome C, and 5 IU/mL catalase in PBS, pH 7.0) or negative control mixture (COX reaction mixture with 2.5 mM sodium azide). Sections were washed twice for 5 min in PBS, then dehydrated for 2 min each in an ethanol series (70%, 70%, 95%, 95%, 99.5%) and cleared in xylenes for 10 min. After air-drying, sections were mounted with Permount (Fisher) under glass coverslips. Brightfield images of sections in a region 100-200 µm from the optic nerve were captured using a Nikon Eclipse E1000 light microscope with a 40X oil objective and 3840 x 3072 pixel resolution. Images were captured using identical lighting conditions, and blindly analyzed using ImageJ software (RRID:SCR_002285). Enzyme activity in single cells was quantified by measuring stain intensity for a circular ROI placed in the center of cone mitochondrial clusters.

**Isotopic $^{13}$C succinate labeling.** Krebs-Ringer bicarbonate (KRB) buffer (98.5 mM NaCl, 4.9 mM KCl, 1.2 mM $KH_2PO_4$ 1.2 mM $MgSO_4$-$7H_2O$, 20 mM HEPES, 2.6 mM $CaCl_2$-$2H_2O$, 25.9 mM $NaHCO_3$) optimized for isotopic labeling experiments in retinas was used for labeling experiments. All reagents for metabolite analyses were acquired from Sigma-Aldrich unless noted. Wild-type zebrafish eyes were dissected at RT (~27ºC) in KRB containing 1 mM each unlabeled glucose and succinate. Retinas were then placed in KRB containing 1 mM D-[U-$^{13}$C]-succinate (Cambridge Isotope Laboratories, CLM-1571) with 1 mM unlabeled glucose and incubated for the specified time points at 28°C (5% $CO_2$ and room oxygen), then washed in PBS and flash frozen in liquid nitrogen.

**Mass spectrometry sample preparation.** Metabolites were extracted from retinas using 150 µL 80% MeOH on dry ice, sonicated, and lyophilized at RT until dry. Extracted metabolites were derivatized in 9 µL of 20 mg/mL methoxyamine HCl dissolved in pyridine at 37ºC for 90 min, and subsequently with 9 µL tertbutyldimethylsilyl-N-methyltrifluoroacetamide at 70ºC for 60 min. Metabolites were analyzed on an Agilent 7890/5975C GC/MS using selected-ion monitoring. Peaks were manually integrated using MSD ChemStation software (Agilent, RRID:SCR_015742), and correction for natural isotope abundance was performed using the software Isocor (Millard et al. 2012).

**Data analysis.** Data were processed using Microsoft Excel (RRID:SCR_016137), and all statistical tests were performed using GraphPad Prism (RRID:SCR_002798). Table 2 lists Ns and statistical analysis methods for each figure. 3-D renderings and animations from SBFSEM were created using Blender (RRID:SCR_008606).
